# Independence of Cued and Contextual Components of Fear Conditioning is Gated by the Lateral Habenula

**DOI:** 10.1101/2020.07.12.197319

**Authors:** Tomas E. Sachella, Marina R. Ihidoype, Christophe D. Proulx, Diego E. Pafundo, Jorge H. Medina, Pablo Mendez, Joaquin Piriz

## Abstract

Fear is an extreme form of aversion that, if inappropriately generalized, initiates pathological conditions such as panic or anxiety. Fear conditioning (FC) is the best understood model of fear learning. During FC two independent associations link the cue and the training context to fear expression. The lateral habenula (LHb) is a general encoder of aversion. However, its role in fear learning has not been intensively studied. Here we studied the role of the LHb in FC using optogenetics and pharmacological tools in rats. Disrupting the neuronal activity of the LHb during training abolishes the expression of fear to isolated presentation of the training context or the cue, yet the recall of both associations when the cue is played in the training context reveals a conserved memory. Our results demonstrate that the LHb is required for the formation of independently expressible contextual and cued memories, a previously uncharacterized role in FC.

## Introduction

Fear is an extreme form of aversion characterized by an uncontrollable reaction to a threatening stimulus. Pavlovian fear conditioning (FC) is probably the most studied and best understood model of fear learning (Maren, 2008; Maren and Quirk, 2004). In FC an association is made between a neutral stimulus and a biologically-relevant aversive stimulus called unconditioned stimulus (US). Most frequently, FC training involves the pairing of a tone, named cue, and an electric foot-shock US. It has long been known that such protocol leads to two independent associations, one relating the cue to the US, and the other linking the context where the training took place to the US (Kim and Fanselow, 1992; Phillips and Ledoux, 1992). Prevailing models postulate that tone-US association takes place in cortical and thalamic auditory inputs to the lateral amygdala (Bocchio et al., 2017; Nabavi et al., 2014; Rumpel et al., 2005; Sah et al., 2008), while context-US association involves context encoding circuits centered in the hippocampus which send contextual representation to the amygdala (Kim and Cho, 2020; Maren et al., 2013).

The lateral habenula (LHb) is a hub for the processing of aversive information in the brain. Aversion-related information reaches the LHb from the basal ganglia and numerous structures of the limbic system including the Central Amygdala (Hu et al., 2020). In turn, the LHb projects to the brain stem, where it is one of the few structures that controls both serotoninergic and dopaminergic systems (Hikosaka, 2010; Hu et al., 2020; Proulx et al., 2014). Notably the LHb and its downstream targets are activated by electric foot-shocks (Lecca et al., 2020; Szőnyi et al., 2019b; Trusel et al., 2019; Wang et al., 2017) and develop responses to cues that predict these aversive stimuli along with the appearance of conditioned responses (Lazaridis et al., 2019; Trusel et al., 2019; Wang et al., 2017). This suggests the LHb may play a role in FC. However, only a few studies examined this possibility (Durieux et al., 2020; Song et al., 2017; Wang et al., 2013). Thus, in this work we investigated the participation of the LHb in the acquisition of contextual and cued FC by means of pharmacological and optogenetics manipulations. We show that interfering with the neuronal activity of the LHb during FC training severely impairs the recall of contextual and cued FC memories. However, when the cue is played in the training context, an intact FC memory is expressed. Thus, our results suggest that the LHb plays a central position in FC learning, affecting not only context and cued FC but also,, remarkably, theirs interaction.

## Results

### Pharmacological inactivation of the LHb impairs contextual fear conditioning

To examine the role of the LHb in contextual FC, we analyzed the effect of its inactivation during training. For that purpose, we performed surgeries in rats, to implant bilateral intracerebral cannulae aimed at the LHb. After a recovery time of 10 to 12 days, animals were bilaterally infused with the GABA-A agonist, muscimol, or vehicle in the LHb and were trained in contextual FC thirty minutes later. During training rats were placed in a FC chamber and presented with four unsignaled mild foot-shocks (Figure 1A). We observed that the infusion of muscimol in the LHb did not modify freezing behavior during FC training (Supp. Fig. 1). To evaluate contextual FC, animals were placed back in the conditioning chamber 7 days after training and freezing behavior was quantified. Muscimol group displayed significantly lower levels of freezing than the control group, suggesting that inactivation of the LHb during contextual FC training impairs long-term memory formation (Figure 1B).

**Figure 1.**
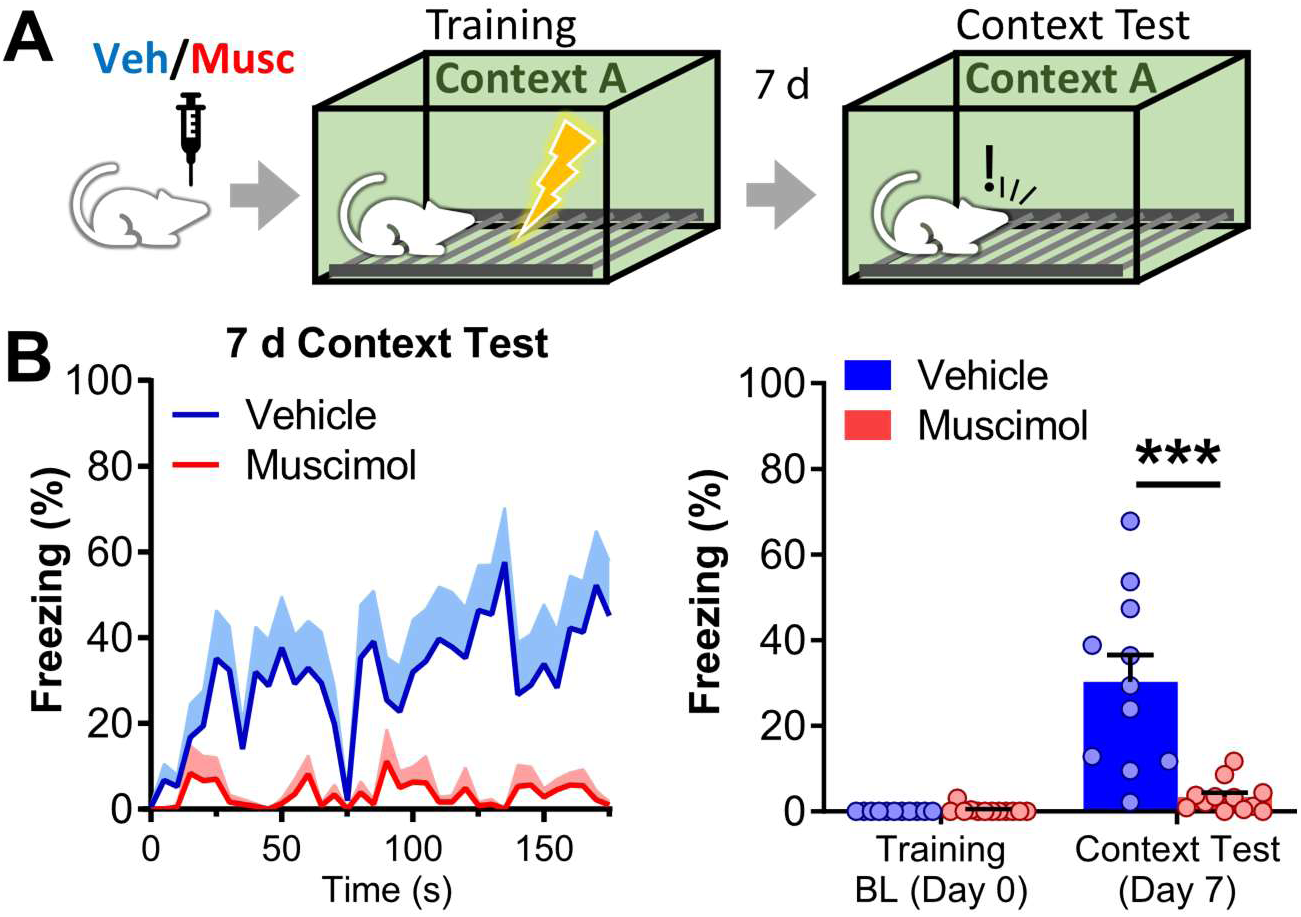
Inactivation of the LHb blocks the formation of contextual FC long-term memory. A) Experiment diagram: bilateral vehicle/muscimol intra LHb infusions were performed 30 minutes before training. Contextual FC was tested 7 days later. During training subjects were let to freely explore the cage for a baseline period of 197 s. After that they were exposed to 4 foot-shocks (0.6 mA, 3 s) interspaced by 87 s. In the test session animals were re-exposed to the training context for 180 s and freezing was quantified. B) Test of contextual FC memory. Left panel: freezing over time. Right panel: average freezing. Freezing in the muscimol group was lower than in the control group. (n_vehicle_ = 11, n_muscimol_ = 12). In freezing over time plot, line represents intersubjects’ mean, and shaded area represents +SEM. In bar plot, each dot represents a subject and bars represent mean and errors bars +SEM. *** p < 0.001. Additional statistics information could be found in the Statistics Details supplementary file.

In a separate cohort of animals, we observed that the infusion of muscimol in the LHb before the exposure to an open field (OF) did not modify locomotion or exploratory behavior (Supp. Fig. 2), indicating that inactivation of the LHb does not have a general effect over mobility or context exploration. Moreover, upon a subsequent re-exposure to the OF 48 hours later, muscimol infused animals displayed a decrease in exploratory behavior equivalent to controls (Supp. Fig. 2). This decrease in exploration is an index of a non-associative habituation memory that depends on context encoding by the hippocampus (Vianna, 2000; Winograd and Viola, 2004). Thus, this observation suggests that the inactivation of the LHb by muscimol does not induce a general deficit in context encoding.

### Pharmacological inactivation of the LHb impairs cued fear conditioning

Having observed that the inactivation of the LHb impairs the formation of contextual FC we extended our experiments analyzing the effect of LHb inactivation on cued FC memory formation. Animals were infused with muscimol or vehicle in the LHb, and 30 minutes later they were trained with cued FC, in which each of the 4 foot-shocks was preceded by a 17-second-long tone (Figure 2A). Cued FC memory was tested 7 days later by exposing the animals to the tone in a novel context (Context B). Behavior during training confirmed that the inactivation of the LHb does not affect the surge of freezing elicited by shock exposure (Supp. Fig. 3). During test, the presentation of the cue elicited a robust freezing in the vehicle group that was markedly reduced in the muscimol group (Figure 2B). These results indicate that cued FC memory is also impaired by the inactivation of the LHb during training. Thus, inactivation of the LHb during FC training impairs both contextual and cued FC long-term memories.

**Figure 2.**
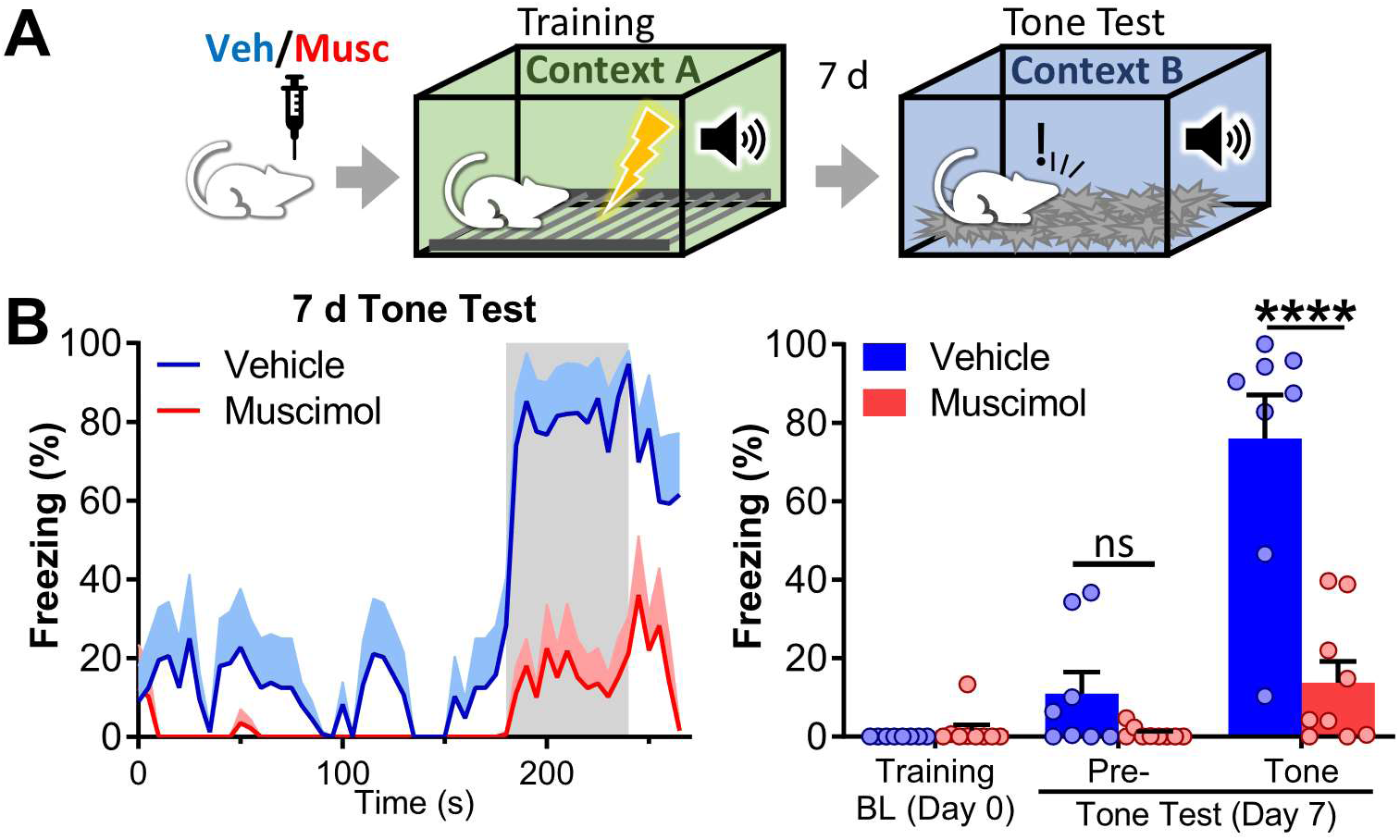
Inactivation of the LHb blocks the formation of cued FC long-term memory. A) Experiment diagram: bilateral vehicle/muscimol intra LHb infusions were performed 30 minutes before training. Cued FC was tested 7 days later. During training, subjects were let to freely explore the cage for a baseline period of 180 s that was followed by 4 tone-shock pairings (17 s of tone followed by 3 s, 0.6 mA shock) interspaced by 70 s. During test, tone was presented for 60 s after a pre-tone period of 180 s. B) Left panel: freezing over time. Gray areas indicate tone presentation. Right panel: average freezing for pre- and tone periods. Freezing during pre-tone period was not different between vehicle and muscimol groups. In contrast, during tone presentation, a highly significant reduction in freezing was observed in the muscimol group. (n_vehicle_ = 8, n_muscimol_ = 9). In freezing over time plot, line represents intersubjects’ mean, and shaded area represents +SEM. In bar plot each dot represents a subject and bars represent mean + SEM. ns p > 0.05, **** p < 0.0001. Additional statistics information could be found in the Statistics Details supplementary file.

Long-term memories tend to decay over time, a phenomenon that could be already evident 7 days after memory formation (Bekinschtein et al., 2007; Rossato et al., 2009). To test whether an accelerated decay could account for deficits of the contextual and cued fear memories, we repeated our previous experiment but tested cued and contextual memories 24 hours after training. Deficits of both memories were also evident at that time point (Supp. Fig. 4), further suggesting that inactivation of the LHb during FC training has a detrimental effect over the formation of long-term contextual and cued FC memories.

As a control for spatial specificity of muscimol infusion we implanted cannulae targeting areas around the LHb and infused muscimol before cued FC training (Supp. Fig. 5A). During tests, performed 7 days later, we found that neither infusion of muscimol 1 mm dorsal, lateral, or ventral to the LHb affected freezing behavior (Supp. Fig. 5B). Moreover, we did not see a reduction in freezing behavior during cued or contextual FC test in muscimol infused animals implanted with cannulae aimed at the LHb in which histological control showed cannulations bilaterally missed the LHb (Supp. Fig. 6).

To control for non-transient effects of muscimol infusion, we trained in cued FC the animals that had previously undergone muscimol inactivation of the LHb before exposure to an OF (Supp. Fig. 2). In these animals, we found no differences in freezing behavior during either cued FC training or testing (Supp. Fig. 7), demonstrating the transient nature of that manipulation. Thus, deficits in contextual and cued FC in the muscimol groups should be attributed to a transient inactivation of the LHb during training.

### Context + tone test reveals a conserved fear conditioning memory

To further investigate the extent to which inactivation of the LHb impairs FC learning, we performed additional experiments where freezing to the cue was evaluated in the training context (Context A), a condition we called context + tone (Figure 3A, Supp. Fig. 8). In that test the pre-tone period is a readout of the contextual component of the memory after a cued FC training. Indeed, during the pre-tone period we observed a clear reduction of freezing in the muscimol group, confirming that the inactivation of the LHb during training impairs contextual FC (Figure 3B). Presenting the tone further increased freezing in the control group and, most notably, it also induced a robust freezing in the muscimol group, which reached freezing values equivalent to the control group (Figure 3B). Thus, under the inactivation of the LHb, a FC memory is formed that could be effectively retrieved when the cue is presented in the conditioning context.

**Figure 3.**
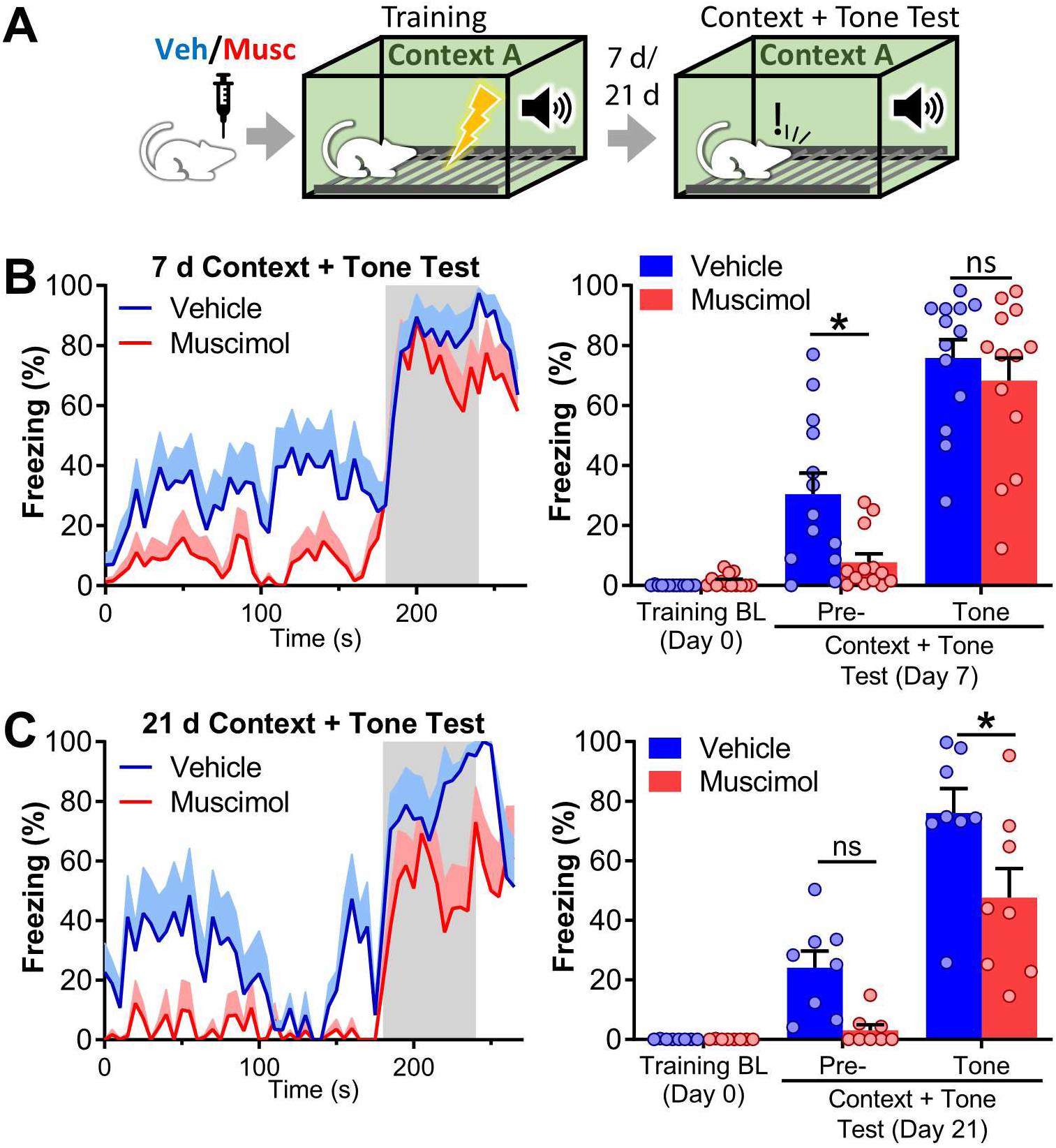
FC memory formed under inactivation of the LHb could be retrieved in context + tone conditions and showed decreased temporal stability. A) Experiment diagram: bilateral intra LHb infusions of vehicle/muscimol were performed 30 minutes before training. Animals were trained in cued FC. Test was performed 7 or 21 days later in the same context used for training (Context A). The tone was presented for 60 s after a pre-tone period of 180 s. B) Freezing during context + tone test 7 days after training. Left panel: freezing over time. Gray area indicates tone presentation. Right panel: average freezing for pre- and tone period. Freezing to the context was higher in vehicle group. However, tone elicited a robust freezing in the muscimol group that reached values equivalent to the vehicle group. n_vehicle_ = 13, n_muscimol_ = 13. C) Freezing during context + tone test 21 days after training. Left panel: freezing over time. Gray area indicates tone presentation. Right panel: average freezing for pre- and tone periods. 21 days after training freezing in context + tone conditions in the muscimol group was lower than in the vehicle group indicating a diminished temporal stability of context + tone memory. n_vehicle_ = 8, n_muscimol_ = 8. In freezing over time plots, line represents intersubjects’ mean, and shaded area represents +SEM. In bar plots each dot represents a subject and bars represent mean +SEM. ns p > 0.05, * p < 0.05. Additional statistics information could be found in the Statistics Details supplementary file.

In a previous work we demonstrated that the inactivation of the LHb impairs long-term stability of the Inhibitory Avoidance memory (Tomaiuolo et al., 2014). To investigate long-term stability of FC memory formed under the inactivation of the LHb, we evaluated, in a new set of animals, freezing in “context + tone” conditions 3 weeks after training (Figure 3A, C). Freezing levels of the muscimol group in the “context + tone” condition was lower than in the vehicle group 3 weeks after training (Figure 3C), showing that FC training under the inactivation of the LHb generates a weak memory that is harder to retrieve and is also temporarily less stable.

### Artificial activation of the LHb impairs cued and contextual fear conditioning

The increase in the neuronal activity of the LHb has been linked to aversion, which is at the core of fear learning (Barker et al., 2017; Lazaridis et al., 2019; Li et al., 2011; Meye et al., 2015; Shabel et al., 2012; Stamatakis and Stuber, 2012). On the other hand, it has been shown that neuronal activity of the LHb is co-modulated with hippocampal theta rhythm (Aizawa et al., 2013; Bertone-Cueto et al., 2020; Goutagny et al., 2013), and it has been postulated that such modulation reflects an influence of the LHb in context encoding at the hippocampus (Baker et al., 2019; Goutagny et al., 2013). Those two seemingly unrelated observations delineate two different scenarios about how the LHb could participate in FC. In the first one, an increase or decrease of neuronal activity of the LHb would lead to changes in aversion and consequently in FC, in the second one the LHb would participate in FC through a dynamically modulated pattern of activity which, if disrupted, would affect FC. To get an insight onto these possibilities we analyzed the impact of sustained excitation of the LHb on FC. For this purpose, we used optogenetics to optically stimulate the LHb during training. We chose an optogenetic over chemogenetic approach since it allows excitation to be limited to the training period, discarding potential effects of the LHb during the consolidation phase. To achieve sustained stimulation of the LHb during training we infused adeno-associated viruses (AAVs) encoding the fast opsin oChIEF (Lin et al., 2009) and implanted optic fibers bilaterally immediately above the LHb (Figure 4A, B). Three weeks later subjects were trained in cued FC under a sustained 20 Hz excitation drive with light pulses of 5 ms (Figure 4A), which has been shown not to induce depression-like symptoms (Yang et al., 2018). Patch-clamp recordings from oChIEF expressing neurons confirmed that responses of LHb neurons to optogenetic stimulation is stable for the duration of the training period (Supp. Fig. 9). We also observed that light stimulation did not modify freezing during FC training (Supp. Fig. 10). During cued memory test, oChIEF stimulated animals showed lower freezing to the tone than the control group (Figure 4C, E). Next day, during “context + tone” test session, oChIEF stimulated animals showed almost no freezing to the context but displayed freezing to the cue, reaching levels higher than in the cued memory test performed the previous day (Figure 4D, E). Thus, the excitation of the LHb during the whole training has a deleterious effect over FC learning mostly equivalent to that of pharmacological inhibition, suggesting that proper encoding of the LHb during FC is required for the formation of independent cued and contextual FC memories.

**Figure 4.**
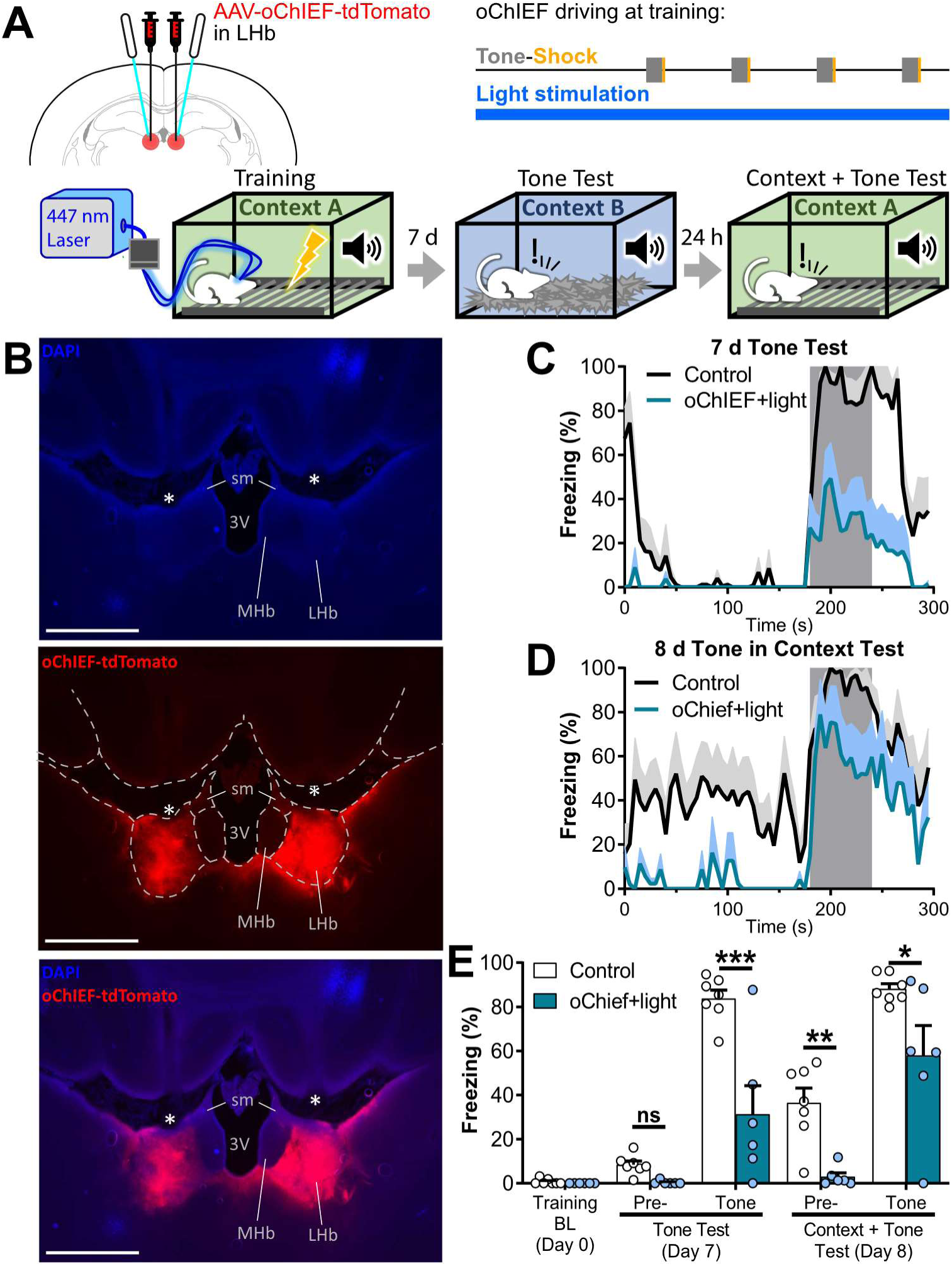
Optogenetic excitation of the LHb during complete FC training impairs contextual and cued FC memories. A) Experiment diagram: Top-Left: animals were bilaterally infected with AAV-oChIEF-tdTomato in the LHb and implanted with optic fibers above the LHb 4 weeks before training. Top-Right: during whole training session the LHb was stimulated with 5 ms light pulses at 20 Hz to disrupt endogenous neuronal activity. Bottom: diagram of training and tests. Cued memory was tested 7 days after training in Context B. The same animals were tested the following day in context + tone conditions to evaluate contextual FC memory and freezing in context + tone condition. B) Microphotographs of the AAV-oChIEF-tdTomato infection (top, middle, bottom: DAPI, tdTomato fluorescence, and merge respectively). Dashed white lines in the middle panel delimitates brain structures. * indicates the optic fiber tract. MHb: medial habenula, sm: stria medullaris, 3V: third ventricle. Scale bars: 1 mm. C, D) Freezing over time during cued test at day 7 (C) and during context + tone test at day 8 (D). Gray area indicates tone presentation. Line represents intersubjects’ mean, and shaded area represents +SEM. E) Average freezing of tone test and context + tone test sessions. Disruption of endogenous LHb activity during training by optogenetic stimulation induced deficits in cued and contextual memories. In bar plots, each dot represents a subject and bars represent mean +SEM. ns p > 0.05, * p < 0.05, ** p < 0.01 *** p < 0.001. n_Control_ = 7, n_oChIEF_ = 6. Additional statistics information could be found in the Statistics Details supplementary file.

### Optogenetic inactivation of the LHb impairs cued fear conditioning

Previous articles have described that neuronal activity at the LHb increases during presentation of the cue and the US in both FC and active avoidance paradigms (Trusel et al., 2019; Wang et al., 2017). We thus tested if inhibiting the LHb during cue and US presentation affects FC. For that purpose, we injected an AAV encoding the inhibitory light-dependent proton pump Archaerhodopsin (ArchT) fused to the fluorescent protein GFP (AAV-CamKIIα-ArchT-GFP) bilaterally in the LHb of rats (Figure 5A). Control rats were injected with AAVs encoding the fluorescent protein GFP (AAV-CamKIIα-GFP) and fiber cannulae were implanted with the tip above the LHb in both groups (Figure 5A, B). Whole cell patch-clamp recordings confirmed that LHb neurons could be inhibited by ArchT (Supp. Fig. 11). Light was applied during the FC training session starting at tone onset and stopping 5 s after shock termination (Figure 5A). We did not observe differences in freezing between ArchT and GFP groups during FC training (Supp. Fig. 12). During cued memory test we found that freezing to the tone was lower in the ArchT than in the GFP group. (Figure 5C, E). On the other hand, in contrast to pharmacological manipulation, contextual memory was not affected by optogenetic inhibition of the LHb during cue and US (Figure 5D, E). Additionally, as observed in pharmacological experiments, context + tone condition evoked similar freezing in ArchT and GFP groups (Figure 5D, E). ArchT mediated inhibition of the LHb for the same amount of time but during inter tone period did not affect cued memory (Supp. Fig. 13). Thus, those results show that the inhibition of the LHb during cue and US presentation is sufficient for the disruption of cued FC but does not affect its contextual component.

**Figure 5.**
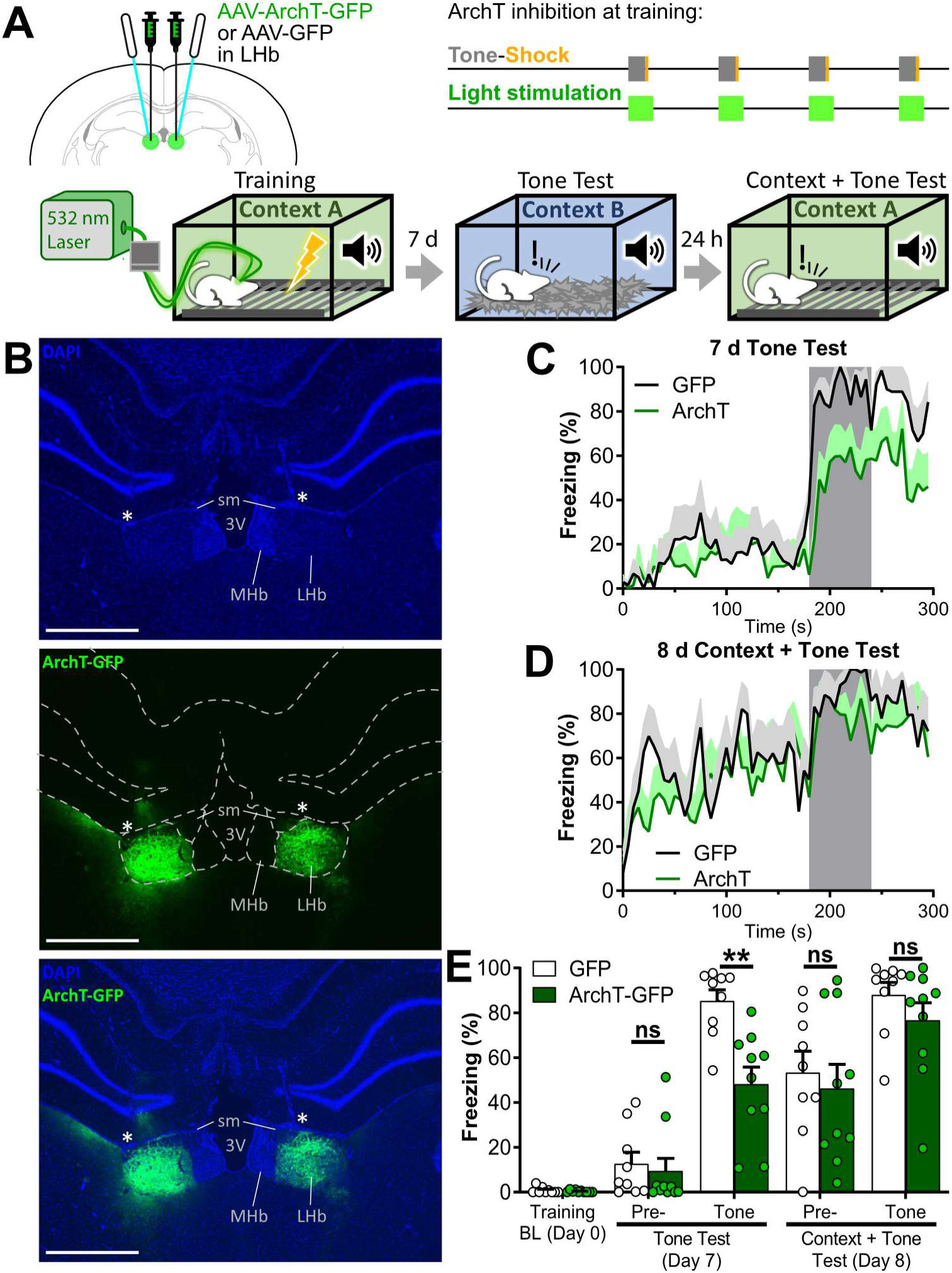
Optogenetic inactivation of the LHb during cue and US, reduced cued but not contextual FC memories. A) Experiment diagram: Top-Left: animals were bilaterally transfected with AAV-ArchT-GFP or AAV-GFP in the LHb and implanted with optic fibers above the LHb 4 weeks before training. Top-Right: during training, optogenetic light stimulation was delivered starting with the tone and stopping 5 seconds after the shock (tone and shock presentations were as previously described for cued FC). Bottom: diagram of training and tests. Cued memory was tested 7 days after training in Context B. The same animals were tested the following day in context + tone conditions to evaluate contextual FC memory and freezing in context + tone condition. B) Microphotographs of the AAV-ArchT-GFP infection (top, middle, bottom: DAPI, GFP, and merge respectively). Dashed white lines in the middle panel delimitates brain structures. * indicates the optic fiber tract. MHb: medial habenula, sm: stria medullaris, 3V: third ventricle. Scale bars: 1 mm. C, D) Freezing over time during tone test at day 7 (C) and during context + tone test at day 8 (D). Gray area indicates tone presentation. Line represents intersubjects’ mean, and shaded area represents +SEM. E) Average freezing on tone test, and context + tone test sessions. During tone test ArchT group displayed lower levels of freezing to the tone than GFP group. The following day, during context + tone test, freezing levels of the ArchT group to both the context and the tone were equivalent to those of the GFP group. In bar plots each dot represents a subject and bars represent mean +SEM. ns p > 0.05, ** p < 0.01. n_GFP_ = 9, n_ArchT_ = 10. Additional statistics information could be found in the Statistics Details supplementary file.

## Discussion

Few previous articles had studied the LHb in the context of FC learning (Barrett and Gonzalez-Lima, 2018; Durieux et al., 2020; Song et al., 2017; Wang et al., 2013). In this work, we directly investigated this subject expanding previous results. We found that disrupting the neuronal activity of the LHb during FC training, either by inhibition, or by sustained optogenetic activation, impairs cue and contextual FC when tested independently. However, presenting the cue in the conditioning context is sufficient to express FC memory, but with a reduced temporal stability. These results suggest that interfering with neuronal activity of the LHb during FC leads to the formation of a weak memory that is harder to retrieve and temporally unstable. In this regard, in a previous article we have shown that the inactivation of the LHb before training in the Inhibitory avoidance generates a temporally less stable memory (Tomaiuolo et al., 2014). Taken together these two observations support the idea that the LHb calibrates strength of aversive memories during acquisition phase.

### Circuits of the LHb related to FC

Considering its connectivity and its role as a general encoder of aversion it is not surprising that the LHb participates in FC. Seminal works by Matsumoto and Hikosaka demonstrated that the LHb encodes punishment and punishment predictive cues (Matsumoto and Hikosaka, 2007) as a phasic increase in activity. More recent works in rodents showed that appearance of LHb responses to cue CSs parallels conditioning in FC (Lazaridis et al., 2019; Wang et al., 2017) and active avoidance (Trusel et al., 2019).

The LHb projects to both dopaminergic and serotoninergic systems both by direct projections as well as indirectly through it projection to the Rostro Medial Tegmental Nucleus (RMTg) (Jhou et al., 2009) and both neuromodulators regulate fear learning (Bissieére et al., 2003; Burghardt and Bauer, 2013; Kwon et al., 2015; Sengupta and Holmes, 2019). In addition, the RMTg itself has been shown to regulate FC (Jhou et al., 2009). Thus, the structures downstream of the LHb participate in FC.

Structures upstream of the LHb have also been implicated in FC learning, most notably the Central Amygdala, which send a dense projection to the LHb (Zhou et al., 2019), and the Medial Prefrontal Cortex (Herry and Johansen, 2014), but also sensory and motivational information relevant to FC reach the LHb from upstream structures such as the Lateral Hypothalamus (Lazaridis et al., 2019; Trusel et al., 2019), the Entopeduncular Nucleus (Li et al., 2019a), the Medial Septum (Pathway et al., 2018), the Median Raphe (Szőnyi et al., 2019b), or the Lateral Preoptic Area (Barker et al., 2017). Thus, the LHb could control FC by gating aversive signal transmission from different sources to serotonergic and dopaminergic centers which ultimately control the formation of aversive memories in its target structures.

### The LHb cued and contextual fear learning

Given the phasic response of the LHb to cue and US (Trusel et al., 2019; Wang et al., 2017) and its role as source of aversive prediction error in monkeys (Matsumoto and Hikosaka, 2009) and rats (Li et al., 2019b), it is conceivable that during cued conditioning the LHb encodes a predictive signal of cue-US contingency as a transient increase in neuronal activity (as postulated also by Li et al., 2019b). Indeed, we found that optogenetic inhibition of the LHb during tone and shock presentation is sufficient to impair cued memory. The importance of the phasic activation of the LHb by the cue and US for cue conditioning could be inferred from our work. It would not be required for the cue-US association to be made but it would regulate the strength of the memory formed. Without proper LHb signaling during training the memory could not be evoked by the cue alone and requires matching contextual information.

Electrophysiological (Congiu et al., 2019; Cui et al., 2018; Lecca et al., 2017; Matsumoto and Hikosaka, 2009, 2007; Yang et al., 2018) and calcium imaging studies (Shabel et al., 2019) showed that 30-50% of the LHb neurons are activated by aversive stimuli such as foot-shock. That subpopulation of LHb neurons activated by foot-shocks would be candidate to be involved in cued conditioning. Recently, transcriptional profiling of LHb neurons identified a specific cluster of neurons in which foot-shocks modulate expression of immediate early genes (Hashikawa et al., 2020). That cluster of LHb cells could be the population identified by electrophysiological recordings. However, this has not been proven yet. Moreover, the lack of a systematic study analyzing the connectivity of foot-shock excited LHb neurons make it impossible to define an LHb centered circuit participating in cued FC with the available information.

In contrast to cued FC, inhibiting LHb during cue and US has no effect on contextual FC, which is instead affected by whole training manipulations of neuronal activity of the LHb (either excitation or inhibition). This discrepancy suggests that the participation of the LHb in contextual conditioning extends beyond its activation by the cue and the US.

It has been proposed elsewhere that the LHb influences context encoding through a functional interaction with the hippocampus (Baker et al., 2019; Goutagny et al., 2013). That hypothesis has received support from physiological data generated by us and other authors showing that firing of the neurons of the LHb is synchronized with hippocampal theta rhythm (Aizawa et al., 2013; Bertone-Cueto et al., 2020; Goutagny et al., 2013). In addition, two recent papers from Nyiri group showed that LHb-innervated Nucleus Incertus (Szőnyi et al., 2019a) and Median Raphe neurons (Szőnyi et al., 2019b) modulate hippocampal theta. Thus, a considerable amount of data supports the hypothesis that the LHb modulates context encoding by the hippocampus. This favors the idea that deficits in contextual FC induced by manipulations of the LHb are consequence of a disturbed encoding of the context. However, we found that, although context by itself does not elicit freezing, it is required for cue evoked freezing, evidencing the retention of contextual information. In addition, inactivation of the LHb does not affect habituation to an OF, which is a form of non-associative learning that is also dependent on contextual representation on the hippocampus (Vianna, 2000; Winograd and Viola, 2004). These two observations contradict the hypothesis that impairment of contextual FC induced by disruption of neuronal activity of the LHb simply represent a deficit in contextual memory. In this regard, the most accepted model of contextual FC postulates that learning take place in two stages (Fanselow, 2010; Rudy et al., 2004), a representative stage in which a cognitive representation of the context is formed in the hippocampus, and a later associative stage in which that contextual representation is associated with the US. The substrates for that association have not been understood. It is tempting to speculate that manipulations of the LHb could help to elucidate them.

### Concluding remarks

Expression of cued and contextual memories have always been considered independent processes. However, our results indicate that such independence depends on LHb signaling during conditioning. Hence, if conditioning takes place without proper activity of the LHb, neither context nor cued memory traces could be independently expressed during test. This observation is novel and assigns the LHb a previously undescribed role within the brain circuits implicated in FC. The circuits implicated in the synergistic action of the context and the cue in fear retrieval are still unknown, thus, their relevance needs to be studied.

Based on our current and previous works we propose that the LHb is a regulator of the aversive memory strength. In FC, the LHb would regulate the strength of both cued and contextual memories probably by different mechanisms. A detailed assessment of that hypothesis could provide valuable information about how the interaction of the hippocampus, the amygdala and the LHb regulates the formation and strength of aversive memories.

## Materials and Methods

### Animals

Experiments were performed in male Wistar rats obtained from the vivariums of the Faculty of Pharmacy and Biochemistry of the Buenos Aires University, Argentina, and Janvier Labs, France. Animals were 5-6 weeks old at the time of surgery. Animals were housed 4 to 6 per cage, with ad libitum access to food and water, under 12-hour light/dark cycle (lights on at 7:00 am), at constant temperature of 22 ± 2 °C. All the procedures were performed during light hours. Experimental procedures were approved by the Animal Care and Use Committee of the University of Buenos Aires (CICUAL), and the Autonomous Community of Madrid (PROEX 167/18).

### Surgeries

#### Pharmacology

rats under deep ketamine/xylazine anesthesia (100 and 5 mg/kg respectively) were bilaterally implanted with 22-Gauge guide cannulae aimed 2.0 mm above the LHb (AP -3.0 mm, ML ±0.7 mm, DV -3.8 mm from Bregma), or 1.0 mm dorsal, ventral or lateral to LHb coordinates in specific experiments. Cannulae were fixed to the skull with three surgical steel screws and dental acrylic. During surgery, animals received a dose of analgesic (meloxicam 0.6 mg/kg) and antibiotic (gentamicin 3 mg/kg). Behavioral procedures began 7-9 days after surgery.

#### Optogenetics

rats under deep isoflurane anesthesia (4 % induction, 1-2 % maintenance, in 0.8 L/minute oxygen) were bilaterally injected with 300 nl of viral vector (AAV8-CamKIIα-ArchT-GFP, 6.2 x 10^12^ viral particles/ml from UNC Vector Core, AAV8-CamKIIα-GFP, 6.3 x 10^12^ viral particles/ml from UNC Vector Core, or AAV8-hSyn-oChIEF-tdTomato homemade as described in (Proulx et al., 2018) per side at the LHb (AP -2.9 mm, ML ± 0.7 mm, DV -5.0 mm from Bregma). For the ArchT experiment, injection of GFP or ArchT-GFP vector was assigned randomly and was balanced across home cages. For the oChIEF experiment, oChIEF-tdTomato vector or saline was assigned randomly and at a 3:1 proportion across home cages. Subsequently, optical fiber implants (200 µm core, 0.45 NA) were bilaterally inserted with the tips aiming just above the LHb, with 4 degrees angle from the sagittal plane (AP -3.0 mm, ML ±1.05 mm, DV -4.5 mm from Bregma), and fixed to the skull with three surgical steel screws and dental acrylic. During surgery, animals received a dose of analgesic (meloxicam 0.6 mg/kg) and antibiotic (gentamicin 3 mg/kg). Behavioral procedures began three weeks after surgery.

#### Electrophysiology

surgery was similar to the optogenetic manipulation experiments with the following differences: 3 weeks old rats were employed, the viral vector injection sites were adjusted according to Bregma-Lambda distance (Bλ) of each rat (AP = -2.9 × Bλ ÷ 8.5 mm, ML = ± 0.7 × Bλ ÷ 8.5 mm from Bregma and DV -4.3 mm from dura mater), no fiber optic implants, nor screws or acrylic was used, and the skin was stitched at the end of surgery.

### Contextual and Cued Fear Conditioning Training

For three consecutive days before the FC training, rats were habituated to handling once a day. There, animals were grasped by hand and slightly restrained against the chest of the investigator for 1-3 minutes, a manipulation similar to the one performed during intracerebral drug infusions. On training day, animals were transferred inside their home cage to a room contiguous to the training room where they acclimatized for 1 hour before the beginning of any other procedure. FC training took place in Context A (electric foot-shock delivering grid floor, black plastic walls and ceiling with one transparent plexiglass side, 50 x 25 x 30 cm wide, width and height, illuminated by white light). Cued FC protocol consisted of a 180 s baseline period of free exploration followed by 4 tone-shock presentations (17 s, 3 kHz, 80 dB tone followed by 3 s, 0.60 mA, square monophasic 60 Hz electric shock) with an inter-stimulus interval of 70 s. Thirty seconds after the last shock, animals were returned to their home cage in the acclimation room. The experimenter who transported the animals between home cages and FC chamber remained blind to the animals’ treatment. When training session ended, home cages were left in the acclimation room for at least one hour before returning them to the animal facility. Ethanol 50 % was used to clean the conditioning cage between subjects. For contextual FC training procedures were equal but the tone was omitted during conditioning. Training was video recorded for posterior analysis.

#### Pharmacology experiments

after the acclimation period, rats were randomly assigned (balanced across home cages) to receive bilateral intra LHb infusions of GABAa agonist, muscimol (Sigma, 60 ng/µl in saline solution, 0.5 µl/side), or saline. Fluorescent green beads were added to both solutions to aid in checking the infusion site (1:1000 dilution of concentrated 1 µm diameter fluorescent green beads, Bangs Laboratories, Indiana, USA). During infusions animals were grasped by hand and slightly restrained in the lap or the arm of the investigator. Infusions were delivered at a rate of 0.5 µl/minute through a 30-Gauge needle extending 2.0 mm beyond the tip of implanted guide cannula and connected to a 10 µl Hamilton syringe by a polyethylene tube. Infusion needle was left in place for an additional minute to minimize backflow. After that animals were returned to their home cage. FC training began 30 minutes later.

#### Optogenetics experiments

after the acclimation period, rats were taken to the FC room and were bilaterally connected to 1 to 2 optic fiber branching patch-cords (200 µm core each, Doric Lenses, Canada). The single end of the branching patch-cord (400 µm core) was connected to a rotary joint (Doric Lenses, Canada) that was attached to the laser source by a simple patch-cord (400 µm core, 0.45 NA, ThorLabs, USA). FC chamber (Context A) had a modification in the ceiling (10 cm hole in the center) to allow optic fiber patch-cord movement. For the ArchT experiment, continuous light of 532 nm at 10 mW at the tip of the fiber implant was delivered by a laser source (CNI, China), starting at tone onset and stopping 5 s after shock termination. For the oChIEF experiment, 5 ms pulses at 20 Hz of 447 nm light at 10 mW at the tip of the fiber implant, were delivered by laser source (Tolket, Argentina) during the whole training. A mixed control group composed of oChIEF injected without light stimulation and saline injected animals with light stimulation was used in that experiment since an AAV-tdTomato was not available. Animals injected with oChIEF on each cage were randomly assigned to light or no-light stimulation. At the end of FC training, patch-cord was disconnected from the fiber implants and animals were returned to home cage.

### Contextual FC Test

The contextual test was performed maintaining the same conditions of the FC training day (e.g. acclimation room, FC chamber lights and investigator’s nitrile gloves) and the experimenter remained blind to the animals’ treatments. During test, animals were placed in the training context (Context A) for 180 s and then returned to its home cage. Test was video recorded for posterior analysis.

### Cued FC Test

Several precautions were taken to avoid generalization during cued test. Home cages were moved to a different acclimation room of the one used in the training day, the investigator used latex gloves, animals were transported between the acclimation room and the test chamber within a plastic box, and acetic acid 1 % m/v was employed for cleaning between subjects. Experimenter remained blind to the animals’ treatments. Test was performed in Context B, a white acrylic box with a transparent plexiglass ceiling (25×25×40 cm wide, width and height) with floor covered in wood shaving and lit up with red light. Cue test consisted of a pre-tone period of 180 s, followed by 60 s tone (same as training tone) and 30 s post-shock period. After that, animals were removed from the test cage and placed back in the home cage. Test was video recorded for posterior analysis. 1 hour after finishing tests, home cages were taken back to the animal facility.

### Context + Tone Test

Context + tone test proceeded as contextual FC test but after 180 s in the test cage the tone was presented for 60 s. 30 s after tone ending animals were returned to their home cage. Experimenter remained blind to the animals’ treatments.

### Open Field

Animals were placed for 15 minutes into an open field (OF; square grey plastic floor and walls 50 cm side x 35 cm height) and video recorded. Experimenter remained blind to the animals’ treatments. Behavior during OF was analyzed offline with ANY-Maze video tracking system (v4.82, Stoelting Co., Wood Dale, USA).

### Histologic Control

Within a week after the end of behavioral procedures animals were euthanized with a lethal dose of ketamine/xylazine, 50 ml of ice-cold phosphate-buffered saline (PBS) 0.1 M was transcardially perfused, followed by 50 ml of paraformaldehyde 4 %. After that, the brain extracted and incubated at 4 °C in paraformaldehyde 4 % overnight. Then, brain was transferred to 30 % m/v sucrose in PBS 0.1 M, incubated for three days at 4 °C and 150 µm coronal sections were cut at a freezing microtome. Sections were mounted with glycerol 50 % v/v in PBS 0.1 M, examined and photographed at 40X magnification with GFP and DAPI filters. For the pharmacological experiments, injection sites were determined by the presence of green beads. Animals in which both infusions were correct were included in the analysis. Animals from the cohorts used in Figure 2B-C in which both infusions were outside the LHb were included in the miss group presented in Figure S3. For the optogenetics experiments, animals with transfection and/or optic fiber misses were not considered in the analysis.

### Freezing analysis

Freezing was manually scored offline. Experimenter was blinded for animaĺs treatment. An Arduino custom system was used for the manual scoring. Briefly, a button was used to synchronize the start of the video and another button was used to register the freezing state while pressed. The Arduino system communicated time and freezing-button state at 10 Hz to the computer, and then saved as CSV file for later analysis. For freezing over time graphs, data was binned in 5 s intervals, and inter-subjectś mean and SEM was calculated.

### Electrophysiological recordings

General methods for slice preparation and recordings were replicated from previous publications of the group (Stefanelli et al., 2016).

### Statistical Analysis

Each animal was taken as an independent measure. During test session of cued or context + tone experiments, average freezing before and during tone presentation were treated as repeated measures. Thus, in most experiments analysis was done using a two-way repeated measures design in which the analyzed factors were “test stage” (pre-tone or tone) and “treatment” (muscimol, optogenetic excitation or inhibition, or the corresponding control group). In experiments in which subjects were tested more than one day, “testing day” was considered an additional factor. Freezing during training session was analyzed separately. There the stage factor comprised freezing during tone or the inter-stimulus period. Statistical analysis of FC experiments was done by generalized linear mixed modelling (GzLMM). Typical distribution of individuals freezing values accumulated at lower and higher values with few individuals freezing around the group mean, indicating a non-normal distribution. We therefore compared models with beta and gaussian distribution (rescaling freezing to y”=[y’(n-1)+0.5]/n as suggested in (Smithson and Verkuilen, 2006). For the model fitting in R (R Foundation for Statistical Computing, Vienna, Austria) we used the ‘glmmTMB’ package (Brooks et al., 2017). Beta distribution provided better adjustment and less tendencies between group residues than gaussian distribution (data not shown). Logit link function was employed. For the GzLMM we considered every interaction between the corresponding fixed factors (test stage, treatment, testing day) and subject was treated as a random factor with random intercepts. ANOVA and contrasts were done applying ‘car’ and ‘emmeans’ packages (Lenth, 2020; Weisberg and Sanford, 2019). Where GzLMM was used. GzLMM ANOVAs used Wald χ^2^ statistics for comparisons since neither likelihood ratio tests nor F tests are supported. Assumptions of homoscedasticity and normal residues were met in the open field experiment; therefore, it was analyzed via conventional two-way repeated measures ANOVA. Each experiment was repeated at least twice in separate cohorts of animals. Each cohort included all the treatments. Sample size calculations were done using power analysis. From pilot context test experiments, we considered a standard deviation of approximately 20 % aiming to detect a minimal difference among groups of 50 %, with an 80 % power, and a type I error of 5 %. Based on these parameters and first assuming gaussian distribution, initially we aimed at an n = 14. Later we readjusted to n = 6 on cued memory tests, where the observed difference between vehicle and muscimol infused groups (aprox. 60 %), standard deviation (aprox. 25 %) and beta distribution at tone tests allowed to maintain about 80 % power and 5 % type I error.

The R scripts generated for the analysis are included as supplemental material.

## Acknowledgements

We would like to thank Dr. Roberto Malinow for his help in the generation of AAV vectors, to Yamila Paez for her technical assistance, and to Sadegh Nabavi, Cecilia Martinez, Mariano Belluscio, Sebastian Giusti and Paula Ospital for their helpful comments. Funding: this work was supported by the following grants: PICT 2016-0034 from the National Agency for Scientific and Technological Promotion of Argentina (ANPCyT) to JM; I-COOP+(CSIC) ref COOPA20198 to PM; NARSAD, Young Investigator Award 2015 (#23861) from the Brain and Behavior Foundation, PICT 2015-2609 and PICT 2017-4594 from the ANPCyT to JP. TS was supported by a predoctoral fellowship from the CONICET. Authors contributions: TES: conceptualization, methodology, software, formal analysis, investigation, data curation, writing-original draft, writing-review & editing, visualization; MRI: validation, formal analysis, investigation, data curation, writing-original draft; CP: validation, pilot experiments, writing-review & editing; DEP: investigation, JHM: resources, writing-review & editing, supervision, funding acquisition; PM: conceptualization, formal analysis, investigation, resources, writing-review & editing, visualization, supervision, funding acquisition; JP: conceptualization, methodology, resources, writing-original draft, writing-review & editing, supervision, project administration, funding acquisition. Competing interests: the authors declare that they have no competing interests. Data and materials availability: all data needed to evaluate the conclusions in the paper are present in the paper and/or the source data files.

---

**Supplementary Figure 1.**
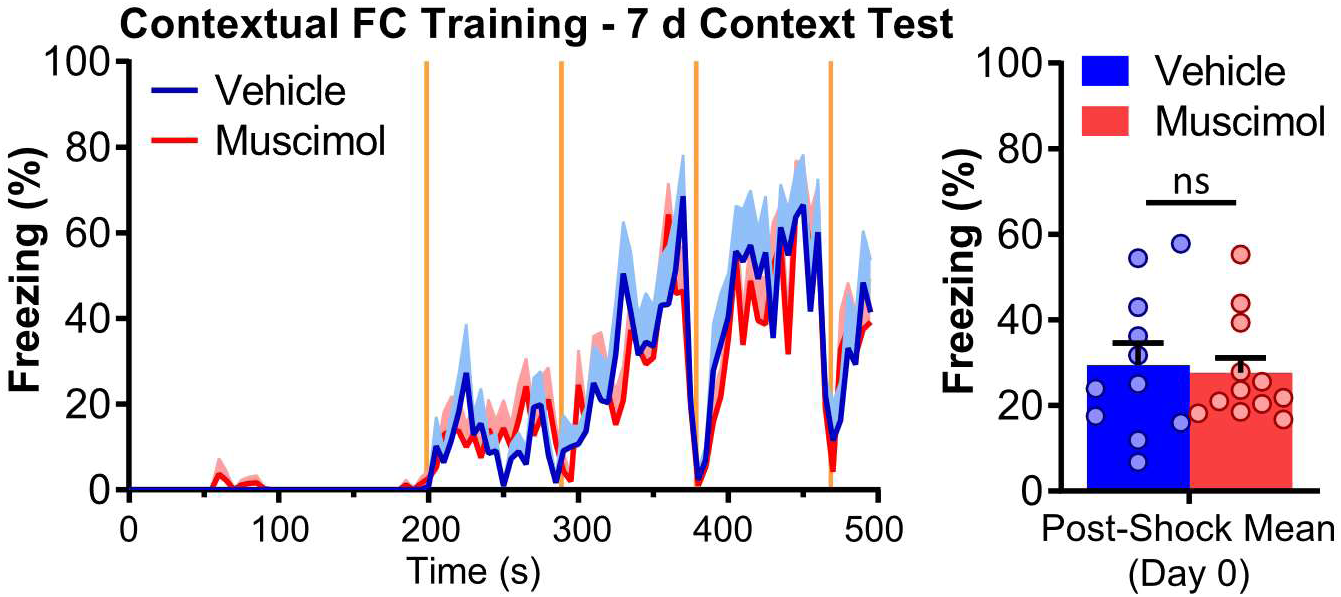
Freezing during training for groups in Figure 1. Left: temporal course of freezing during training. Right: average freezing. No significant differences were observed between muscimol and vehicle group. In freezing over time plot orange shaded areas indicate shock presentation, line plot represent intersubjects’ mean and shaded area over line plot represents +SEM. In bar plot each dot represents a subject and bars represent mean +SEM. ns p > 0.05. n_vehicle_ = 11, n_muscimol_ = 12. Additional statistics information could be found in the Statistics Details supplementary file.

**Supplementary Figure 2.**
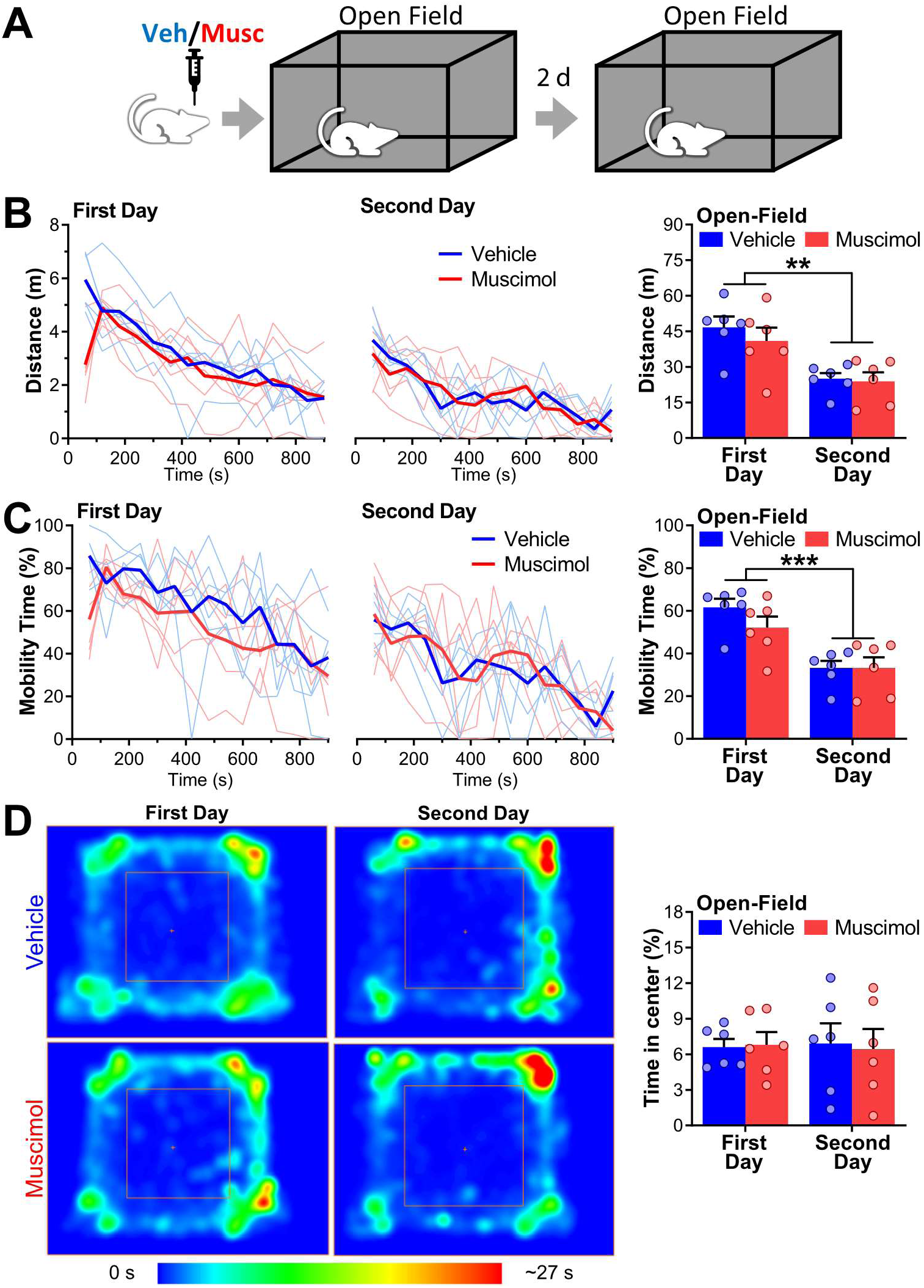
Inactivation of the LHb does not affect locomotion, exploration, or habituation to an Open Field. **A)** Experiment diagram: bilateral intra LHb infusions of vehicle/muscimol were performed 30 minutes before exposing the animals to an OF for 15 minutes. 48 hours later animals were exposed to the same OF for a second time. **B)** Distance traveled over time (60 s bins) during the first day (left) and the second day (right) in the OF. Thin lines represent individuals, thick lines represent the average. Right: total distance traveled. Distance traveled by both groups diminished the second day. No differences between groups were observed. **C)** Percentage of mobility time in the Open Field in each day. Left: percentage of mobility over time (60 s bins) during the first day (left) and the second day (right). Thin lines represent individuals, thick lines represent the average. Right: percentage of mobility time during whole OF assay. Mobility of both groups diminished the second day. No differences between groups were observed. **D)** Heat map representation of spatial occupancy during OF. Left: average heat maps vehicle (top) and muscimol (bottom) groups on the first day (left), and the second day (right). The center zone is indicated by the orange square. Right: percentage of time in the center zone in both groups at each day. Time in the center did not change between days and was equivalent among groups. n_vehicle_ = 6, n_muscimol_ = 6. ** p < 0.01, *** p < 0.001. Additional statistics information could be found in the Statistics Details supplementary file.

**Supplementary Figure 3.**
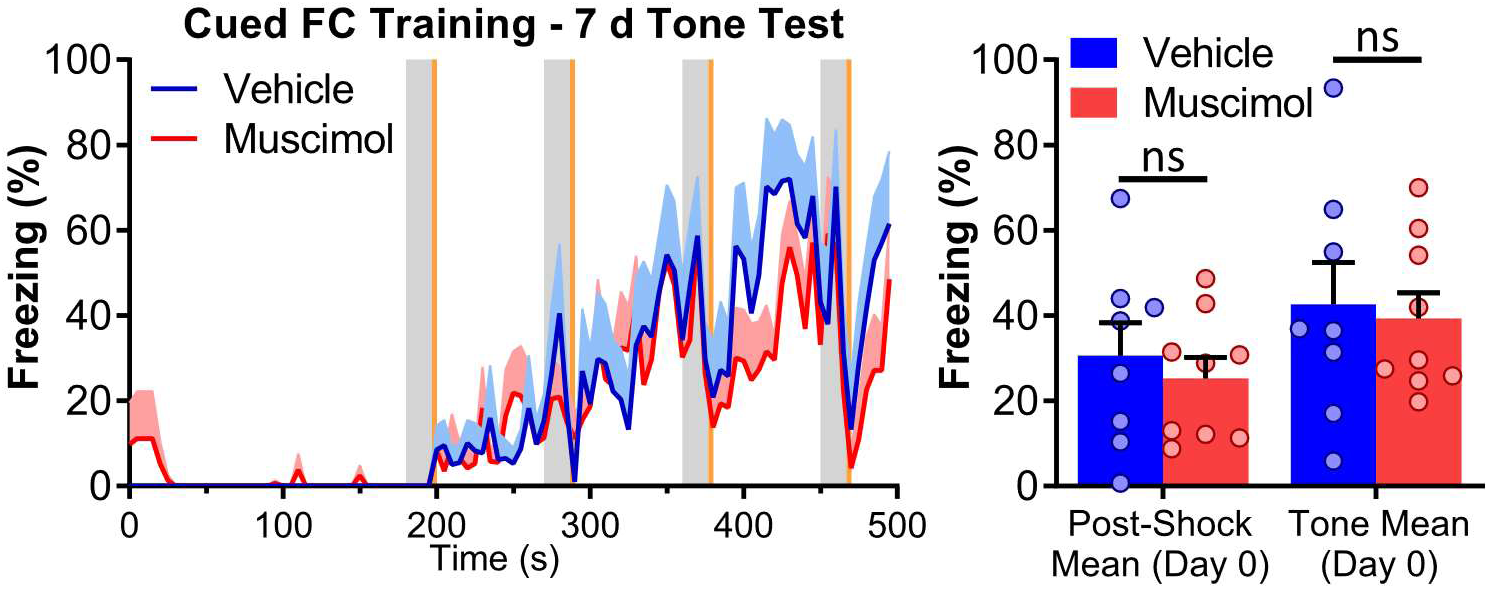
Freezing during training for groups in Figure 2. Left: temporal course of freezing during cued FC training. Right: average freezing between stimuli and during tone presentation. In either condition no significant differences were observed between muscimol and vehicle group. In freezing over time plots gray and orange shaded areas indicate tone and shock presentation respectively, line plots represent intersubjects’ mean and shaded area over line plots represents +SEM. In bar plots each dot represents a subject and bars represent mean +SEM. ns p > 0.05, n_vehicle_ = 8, n_muscimol_ = 9. Additional statistics information could be found in the Statistics Details supplementary file.

**Supplementary Figure 4.**
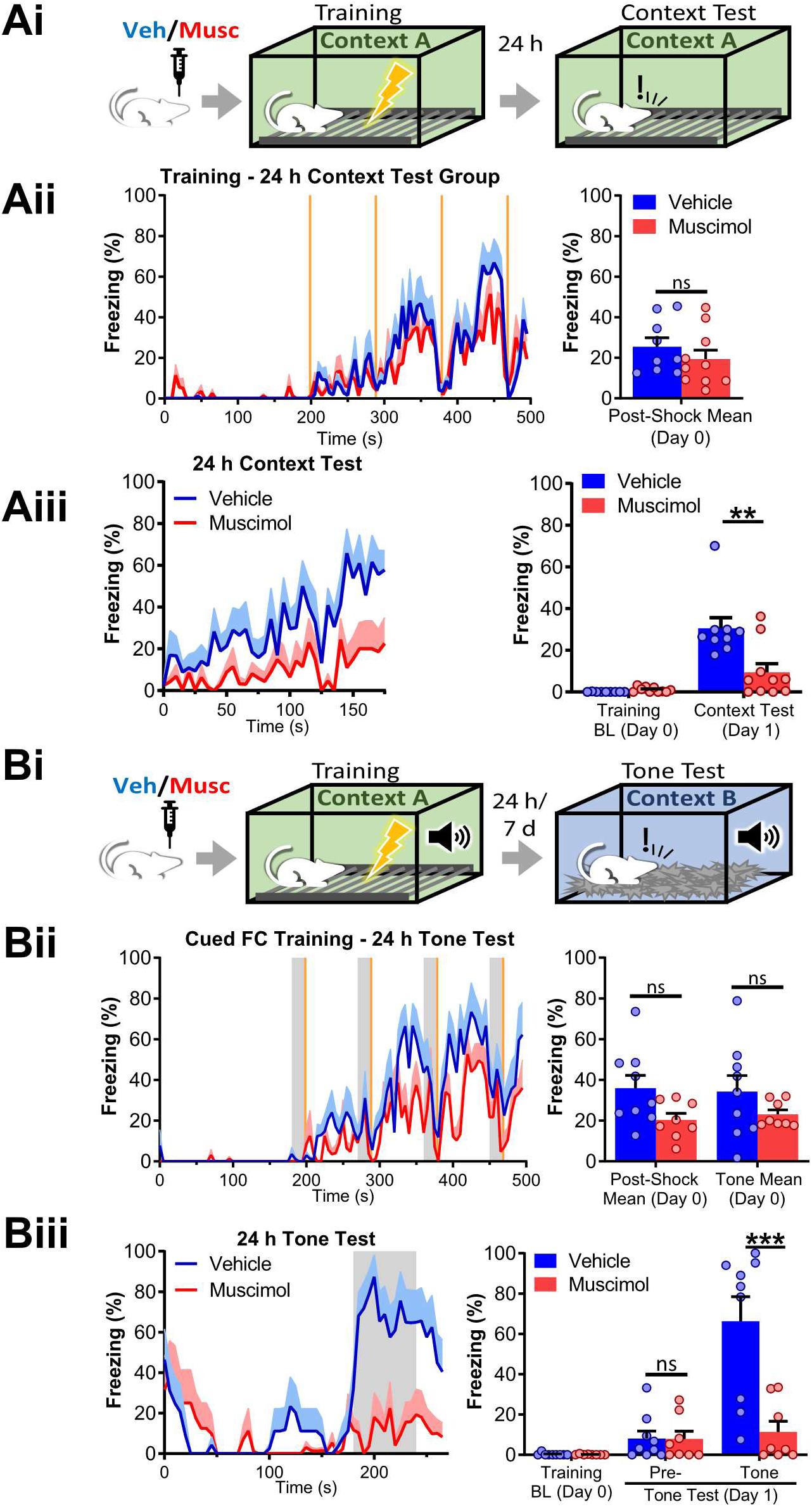
Inactivation of the LHb during training impairs contextual and cued FC memories evaluated 24 hours after training. **A)** Inactivation of the LHb during training impairs 24 h contextual FC memory. i) Experiment diagram: bilateral vehicle/muscimol intra LHb infusions were performed 30 minutes before training contextual fear memory was evaluated 24 hours after training. ii) Left: temporal course of freezing during training. Right: average freezing. No significant differences were observed between muscimol and vehicle groups. iii) 24 hours test of contextual fear memory. Left panel: freezing over time. Right panel: average freezing. Freezing in the muscimol group was lower than in the control group (n_vehicle_ = 9, n_muscimol_ = 10). **B)** Inactivation of the LHb during training impairs 24 h cued FC memory. i) Experiment diagram: bilateral vehicle/muscimol intra LHb infusions were performed 30 minutes before training. Cued FC was tested 24 hour later. ii) Left: temporal course of freezing during cued FC training. Right: average freezing between stimuli and during tone presentation. No significant differences were observed between muscimol and vehicle groups. iii) 24 hours cued FC test. Left panel: freezing over time. Gray areas indicate tone presentation. Right panel: average freezing for pre- and tone period. Freezing during pre-tone period was not different between vehicle and muscimol groups. In contrast, during tone presentation, a highly significant reduction in freezing was observed in the muscimol group (n_vehicle_ = 9, n_muscimol_ = 8). In freezing over time plots, line represents intersubjects’ mean, and shaded area represents +SEM. In bar plots each dot represents a subject and bars represent mean +SEM. ns p> 0.05, *** p < 0.001, **** p < 0.0001. Additional statistics information could be found in the Statistics Details supplementary file.

**Supplementary Figure 5.**
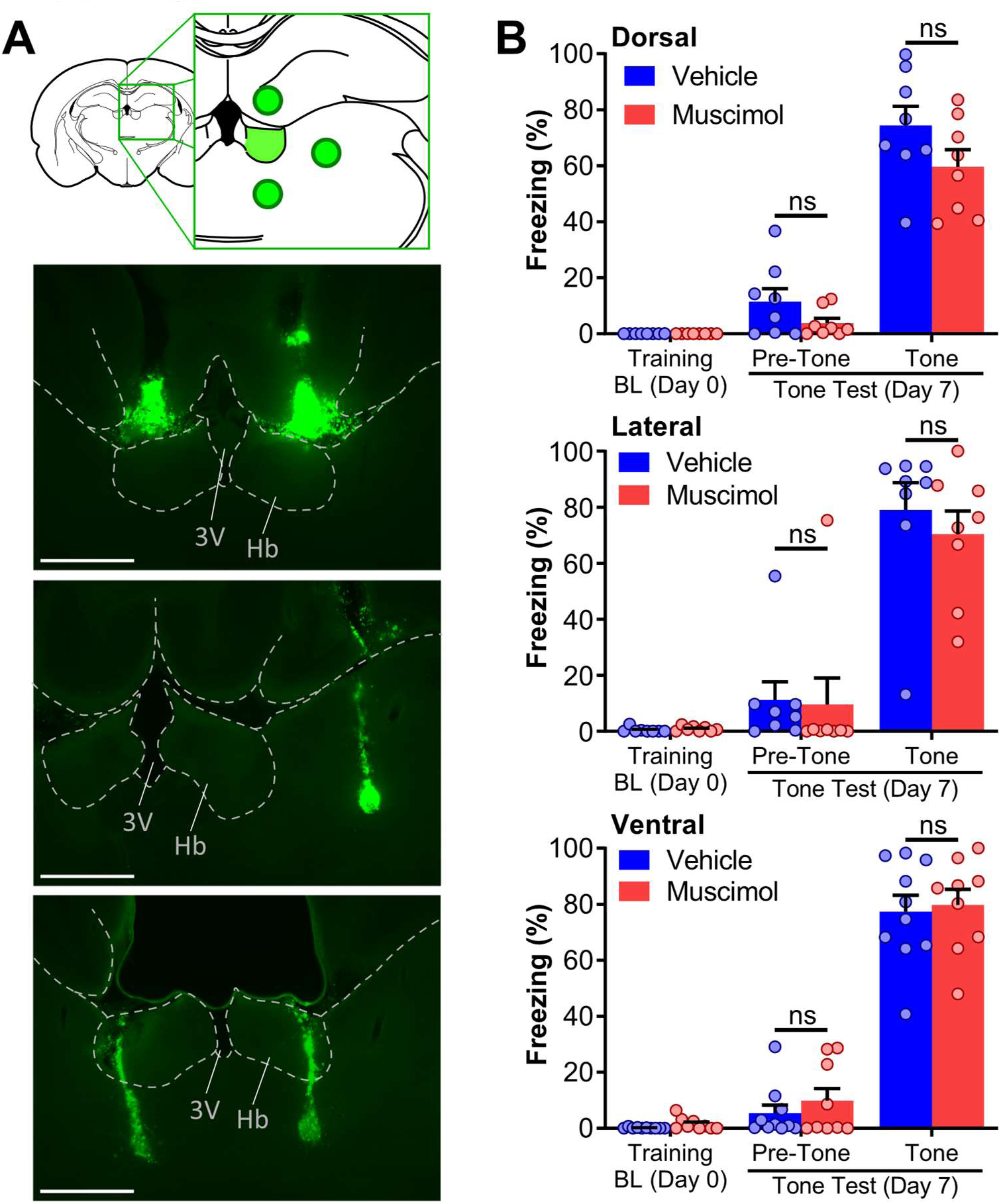
Controls of specificity of muscimol inactivation of the LHb. **A)** Diagram and representative photomicrographs of controls of specificity of muscimol infusion. Infusions were aimed 1 mm dorsal (top picture), lateral (middle picture) or ventral (bottom picture). References: Hb: habenula; 3V: third ventricle. **B)** Freezing during cued memory testing in dorsal, lateral, and ventral controls. Dorsal, ventral, and lateral groups were done as separate experiments. As in Figure 2, infusions were done 30 minutes before training and test was performed 7 days later. Muscimol infusion dorsal, ventral, or lateral to the LHb did not affect tone evoked freezing. In bar plots each dot represents a subject and bars represent mean + SEM. ns p > 0.05. For Dorsal experiment: n_vehicle_ = 8, n_muscimol_ = 8. For Lateral experiment: n_vehicle_ = 8, n_muscimol_ = 8. For Ventral experiment: n_vehicle_ = 10, n_muscimol_ = 9. Additional statistics information could be found in the Statistics Details supplementary file.

**Supplementary Figure 6.**
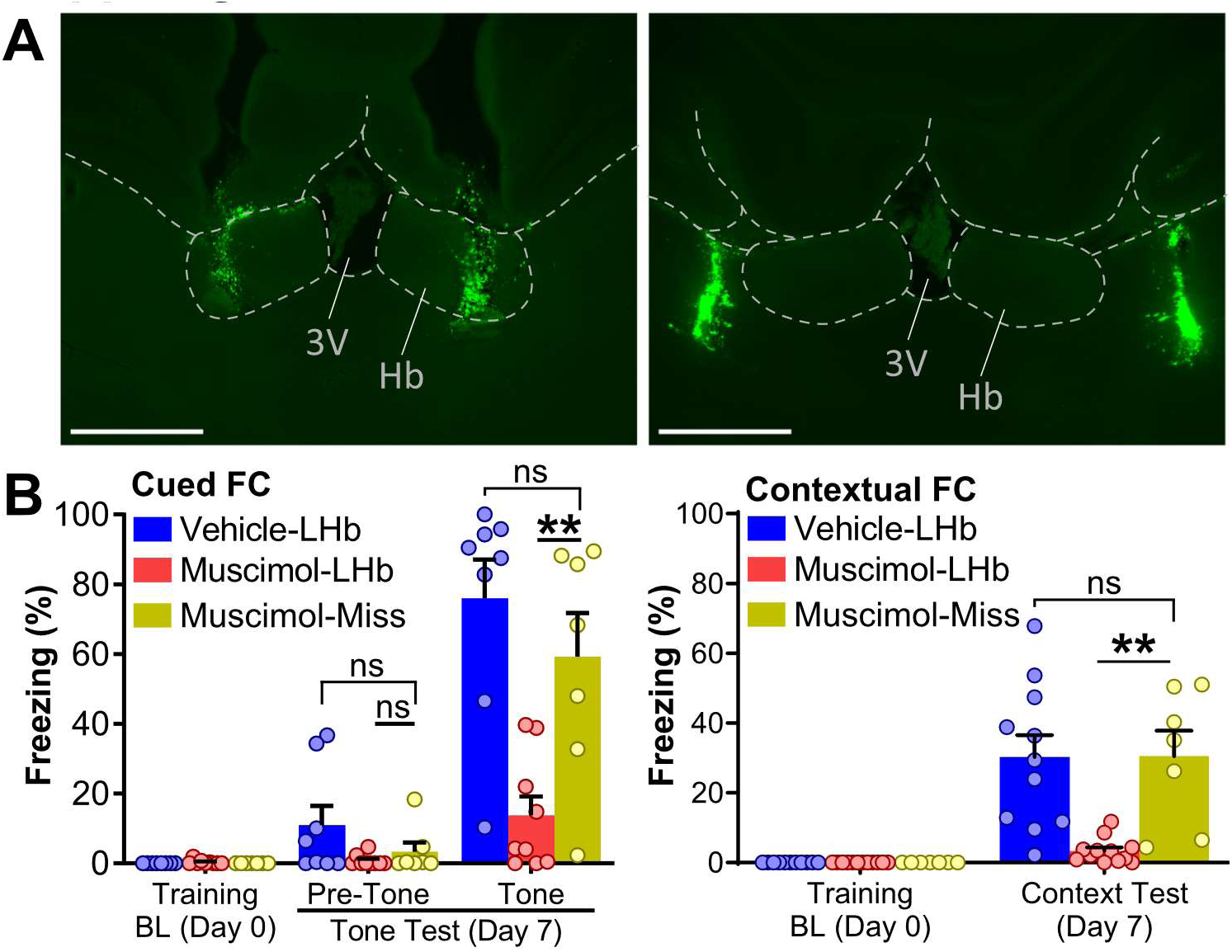
Infusion of muscimol in non-voluntarily bilateral miss-cannulated animals does not block cued or contextual FC. **A)** Photomicrographs of green beads fluorescence of a representative infusion on the LHb (left) or a double miss infusion (right). References: Hb: habenula; 3V: third ventricle. Scale bars: 1 mm. **B)** Freezing during cued (left) or contextual (right) memory testing in miss-LHb groups. Miss-LHb groups consists of animals infused with muscimol in which the two cannulae did not hit the LHb from the same cohorts included in Figures 1B and 2B. As in Figures 1 and 2, infusions were done 30 minutes before training and test was performed 7 days later. At cued memory test (left) Miss-LHb group showed higher freezing to the tone than muscimol-LHb group and equal freezing to the tone than the vehicle-LHb group. The profile was equivalent at the contextual memory test (right), with Miss-LHb and vehicle-LHb groups freezing equally to the tone and each of them higher than the muscimol-LHb group. In bar plots each dot represents a subject and bars represent mean + SEM. Muscimol-LHb and vehicle-LHb groups are taken from Figures 1B and 2B. ns p > 0.05, ** p < 0.01. For cued FC experiment: n_vehicle_ = 8, n_muscimol_ = 9, n_miss_ = 7. For contextual FC experiment: n_vehicle_ = 11, n_muscimol_ = 12, n_miss_ = 7. Additional statistics information could be found in the Statistics Details supplementary file.

**Supplementary Figure 7.**
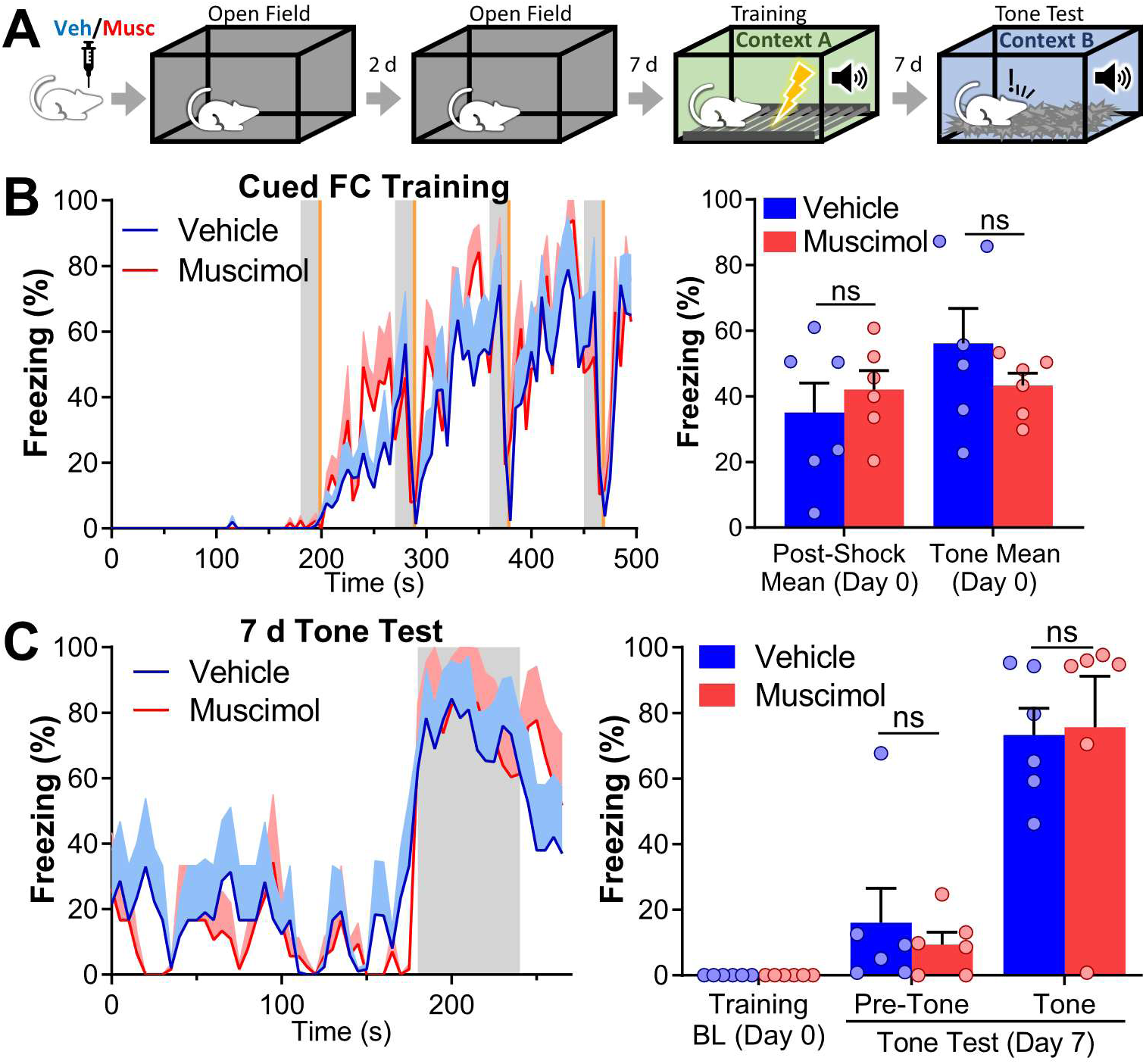
Inactivation of the LHb does not permanently block FC learning. **A)** Experiment diagram: bilateral intra LHb infusions of vehicle/muscimol were performed in the animals that went through the OF (Supplementary Figure 2). Seven days after the second exposure to the OF animals were trained in cued FC. Memory was tested 7 days later. **B)** Cued FC training of the animals that were previously exposed to the OF. Left panel: freezing over time. Tones were presented at times indicated by the gray shaded areas. Shocks were presented at times indicated by the orange shaded areas. Right panel: mean freezing during tone and post-shock period. Freezing in both groups was equivalent. **C)** Test of cued FC memory on the same animals. Left panel: freezing over time. Gray area indicates tone presentation. Right panels: average freezing for pre- and tone period. Freezing was not different between vehicle and muscimol groups. In freezing over time plots, line represents intersubjects’ mean and shaded area represents +SEM. In bar plots each dot represents a subject and bars represent mean + SEM. ns p > 0.05. n_vehicle_ = 6, n_muscimol_ = 6. Additional statistics information could be found in the Statistics Details supplementary file.

**Supplementary Figure 8.**
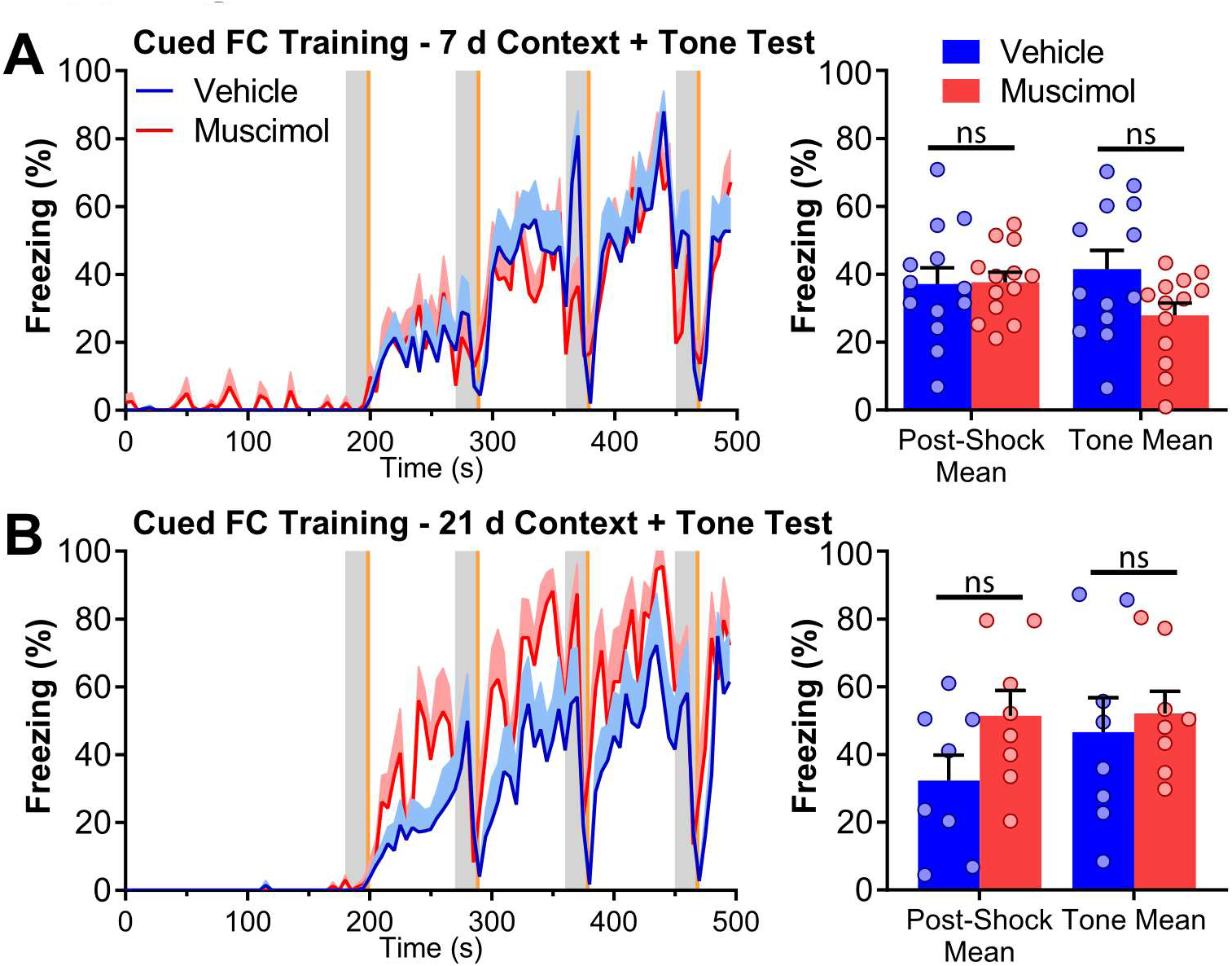
Freezing during training for groups in Figure 3. **A, B)** Left: temporal course of freezing during training for the experiments tested 7 (A) or 21 (B) days after training. Right: average freezing of those experiments. In either condition no significant differences were observed between muscimol and vehicle group. In freezing over time plots orange shaded areas indicate shock presentation, gray shaded areas indicate tone presentation, line plots represent intersubjects’ mean and shaded area over line plots represents +SEM. In bar plots each dot represents a subject and bars represent mean + SEM. ns p > 0.05. For 7 days test experiment: n_vehicle_ = 13, n_muscimol_ = 13. For 21 days test experiment: n_vehicle_ = 8, n_muscimol_ = 8. Additional statistics information could be found in the Statistics Details supplementary file.

**Supplementary Figure 9.**
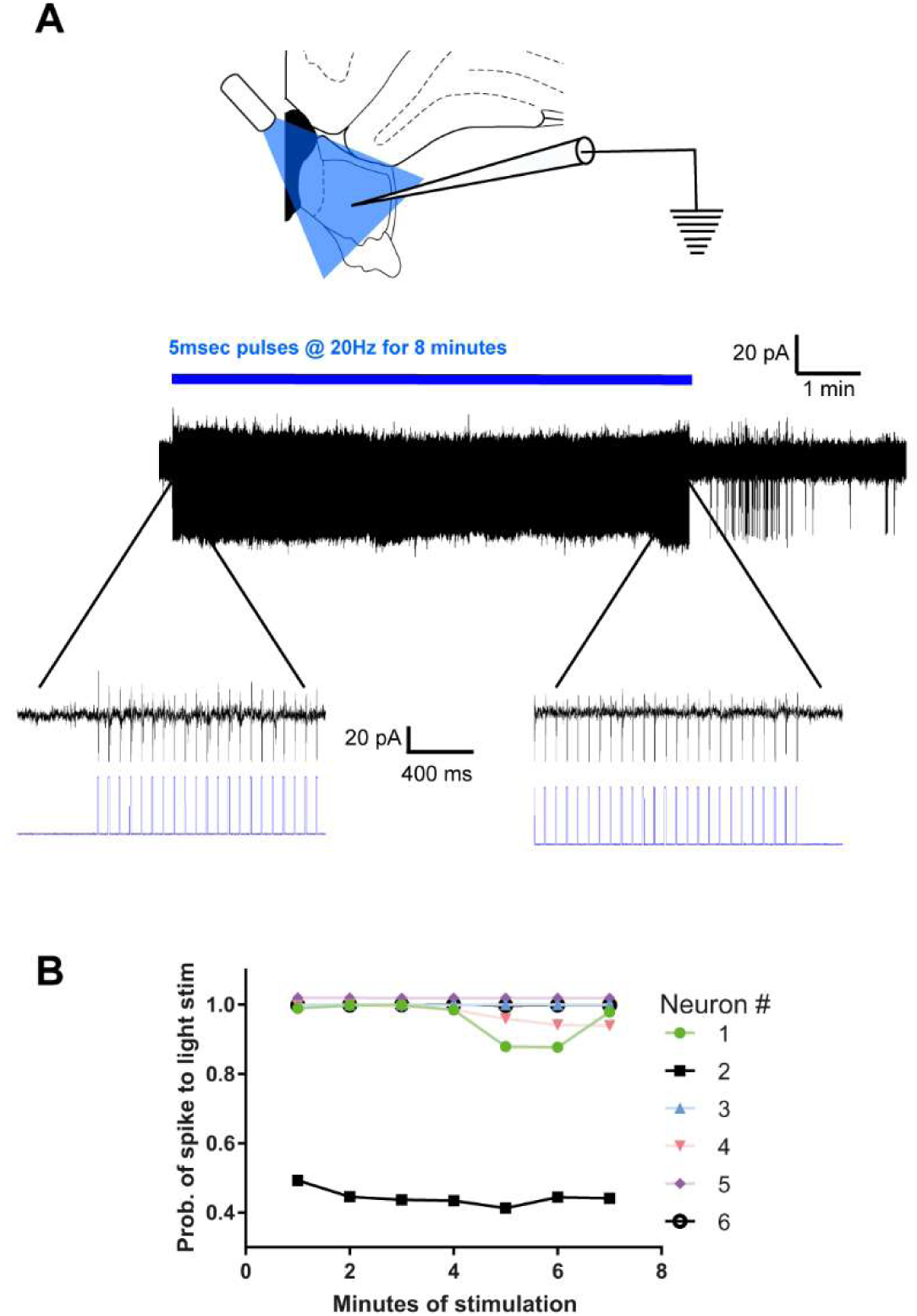
Sustained optogenetic excitation of LHb neurons. **A)** Top, experimental diagram. Neurons of the LHb were recorded in cell-attach configuration and optogenetically stimulated mimicking in vivo stimulation during FC training. Bottom, example of neuron recorded using that protocol. Left and right zoom-in traces show the beginning and the end of the stimulation. **B)** Quantification of all neurons recorded. The graph shows the mean probability of activation by light pulses over a minute of stimulation. The probability of activation did not decrease over time indicating that LHb neurons could sustain responses to optogenetic stimulation for the duration of our behavior protocol.

**Supplementary Figure 10.**
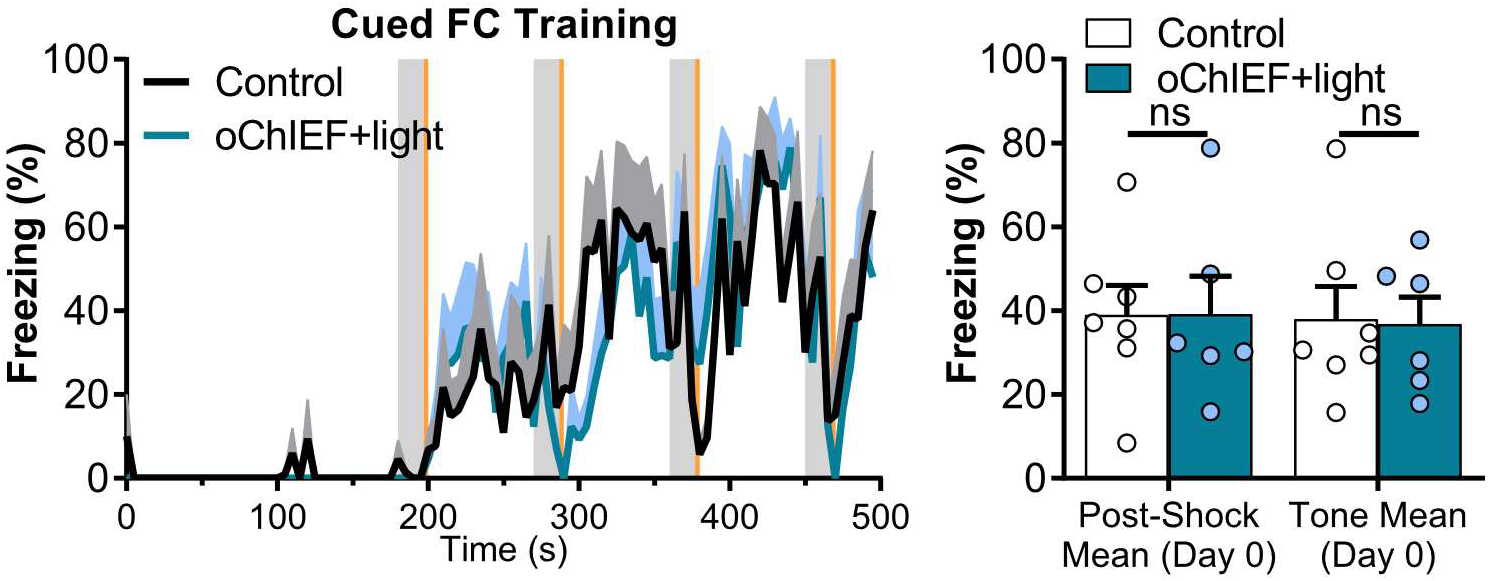
Freezing during training for groups in Figure 4. Left: temporal course of freezing during training in the optogenetics stimulation experiment. Right: average freezing on the same experiment. In freezing over time plot vertical orange shaded areas indicate shock presentation, gray shaded areas indicate tone presentation, line plots represent intersubjects’ mean and shaded area over line plots represents +SEM. No significant differences were observed between control and optogenetically stimulated groups. ns p > 0.05. n_Control_ = 7, n_oChIEF_ = 6. Additional statistics information could be found in the Statistics Details supplementary file.

**Supplementary Figure 11.**
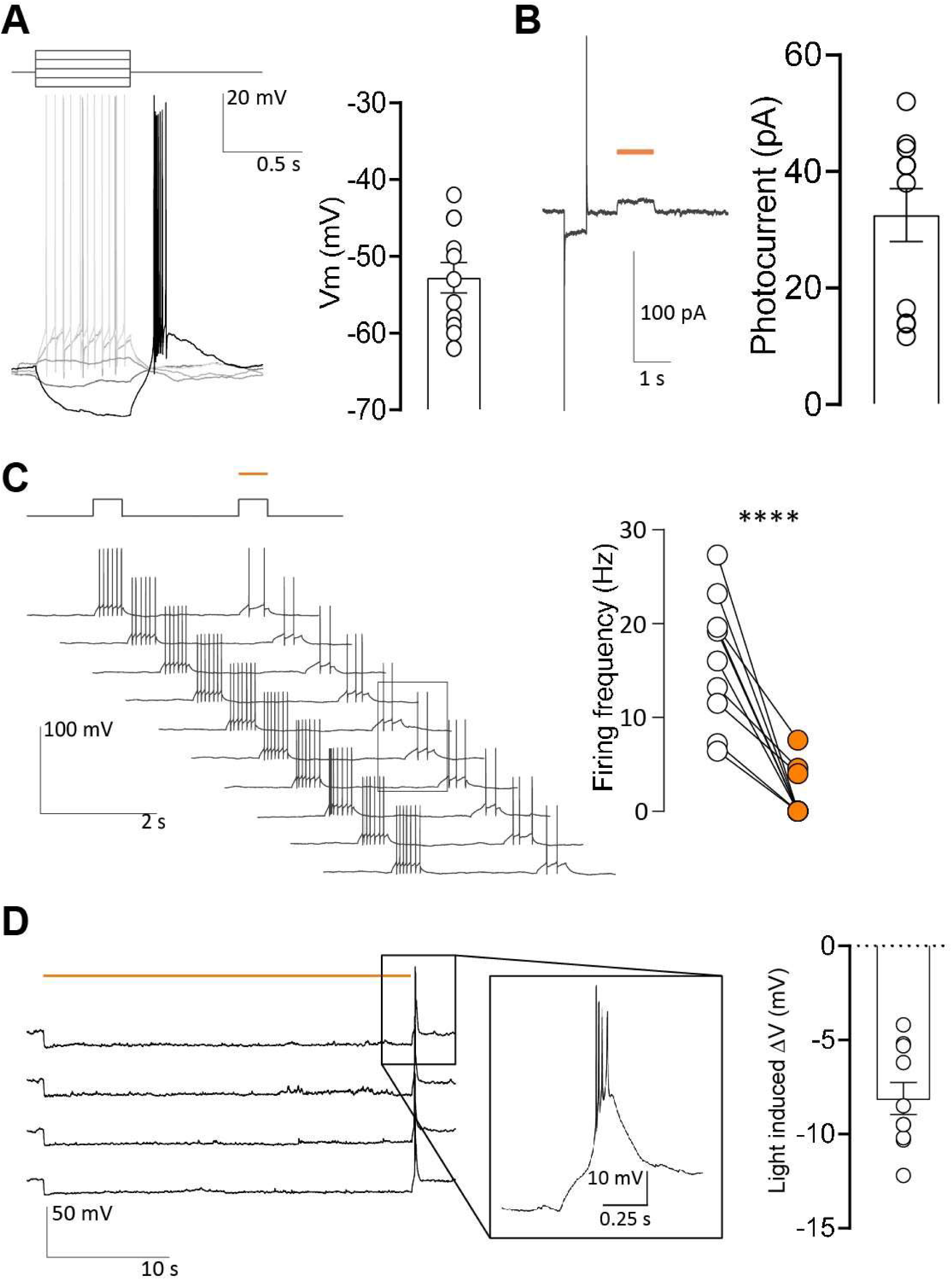
Optogenetic inhibition of LHb neurons by ArchT. **A)** Example of membrane potential response to 600 ms current injection of an ArchT expressing LHb neuron (-40, -10, 15, and 40 pA steps). Rebound spiking was observed after cessation of large hyperpolarizing currents. Right graph: resting membrane potential of recorded neurons; n = 13 neurons from 3 rats. **B)** Light induced photocurrent. ArchT was activated with 590 nm LED for 1 s (orange line) with a nominal power of 0.15 mW; n = 11 neurons from 3 rats. **C)** Light induced action potential suppression in ArchT expressing LHb neurons. Light pulse (590 nm, orange line) was applied every second time neurons were depolarized with positive current injection. Left, representative recordings. Right, data summary. Light activation of ArchT significantly decreases the number of action potentials evoked by current steps in LHb neurons (paired t test, t_(9)_ = 6.25, p < 0.001); n = 9 neurons from 3 rats. **D)** Membrane potential response to a 30 s light pulse (orange line) in LHb neurons. Rebound spiking was observed after termination of light pulse, a representative trace is expanded in the middle traces. This behavior confirms the reported rebound responses of the LHb neurons following long periods of optogenetic inhibition reported previously (Yang et al., 2018). Right: quantification of the stationary light induced ΔVm during the last 5 s of the 30 s pulse; n = 10 neurons from 3 rats.

**Supplementary Figure 12.**
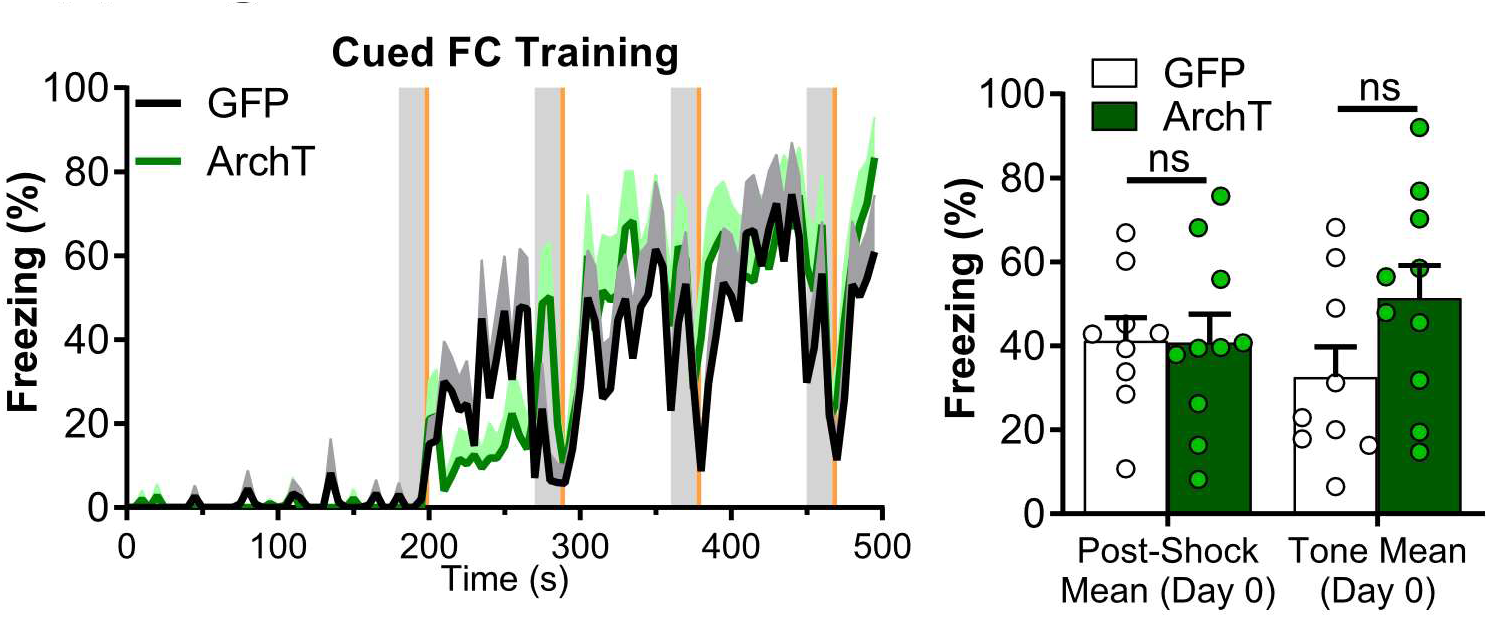
Freezing during training for groups in Figure 5. Left: temporal course of freezing during training in the experiments of optogenetic inhibition with ArchT. Right: average freezing on the same experiment. In freezing over time plot vertical orange shaded areas indicate shock presentation, gray shaded areas indicate tone presentation, line plots represent intersubjects’ mean and shaded area over line plots represents +SEM. No significant differences were observed between control and optogenetically stimulated groups. ns p > 0.05. n_GFP_ = 9, n_ArchT_ = 10. Additional statistics information could be found in the Statistics Details supplementary file.

**Supplementary Figure 13.**
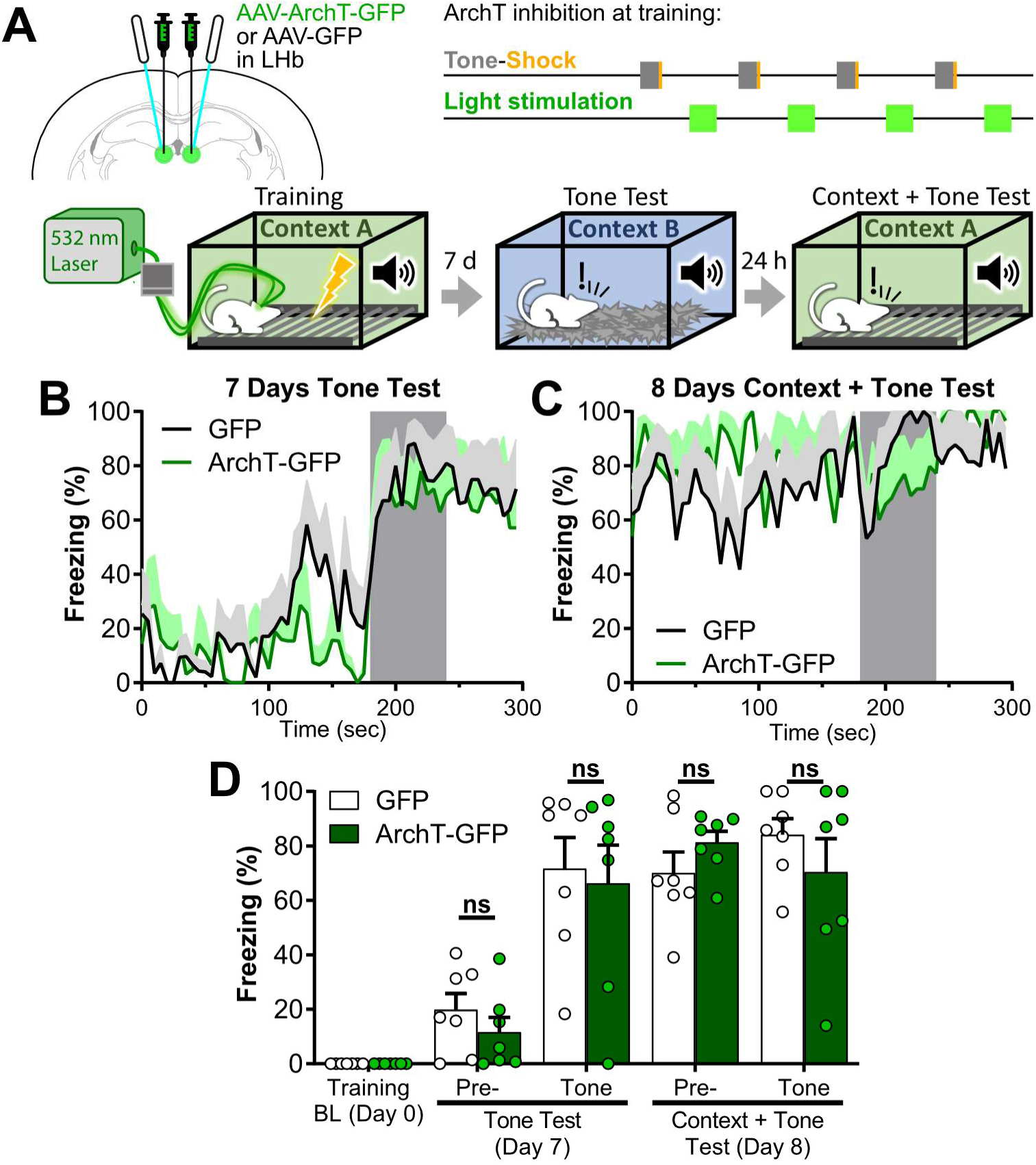
Optogenetic inactivation of the LHb during inter stimulus period, did not affect cued nor contextual FC memories. **A)** Experiment diagram: Top-Left: animals were bilaterally transfected with AAV-ArchT-GFP or AAV-GFP in the LHb and implanted with optic fibers above the LHb 4 weeks before training. Top-Right: during training, optogenetic light stimulation was delivered for 25 seconds during inter stimulus period, starting 30 seconds after each shock termination (tone and shock presentations were as previously described for cued FC). Bottom: diagram of training and tests. Cued memory was tested in Context B 7 days after training. The same animals were tested the following day in context + tone conditions to evaluate contextual FC memory and freezing in context + tone conditions. **B, C)** Freezing over time during tone test at day 7 (B) and during context + tone test at day 8 (C). Gray area indicates tone presentation. Line represents intersubjects’ mean, and shaded area represents +SEM. **D)** Average freezing on tone test, and context + tone test sessions. ArchT-GFP and GFP groups had equivalent freezing over tone test and context + tone test days at each pre-tone and tone phases. In bar plots each dot represents a subject and bars represent mean +SEM. ns p > 0.05. nGFP = 7, nArchT = 7. Additional statistics information could be found in the Statistics Details supplementary file.

## Statistics details

### Figure 1. Statistics

**B)** GzLMM t test for 7 days context test: t_(20)_ = 4.294, p = 0.0004, n_vehicle_ = 11, n_muscimol_ = 12.

### Figure 2. Statistics

**B)** GzLMM Two-way repeated measures ANOVA of treatment, test stage, and interaction for 7 days tone test: χ^2^ treatment _(1)_ = 2.157, p = 0.1420; χ^2^ stage _(1)_ = 19.100, p < 0.0001; χ^2^ interaction _(1)_ = 32.939, p < 0.0001. Sidak’s multiple comparisons test: Pre-tone freezing: vehicle vs. muscimol: t_(28)_ = 1.469, p = 0.2828; Tone freezing: vehicle vs. muscimol: t_(28)_ = 5.396, p < 0.0001; n_vehicle_ = 8, n_muscimol_ = 9.

### Figure 3. Statistics

**B)** GzLMM two-way repeated measures ANOVA of treatment, test stage, and interaction for 7 d context + tone test: χ^2^ treatment _(1)_ = 6.830, p = 0.0090; χ^2^ stage _(1)_ = 44.992, p < 0.0001; χ^2^ interaction _(1)_ = 3.161, p = 0.0754. Sidak’s multiple comparisons test: Pre-tone freezing: vehicle vs. muscimol: t_(46)_ = 2.613, p = 0.0474; Tone freezing: vehicle vs. muscimol: t_(46)_ = 0.899, p = 0.8460; Vehicle: pre-tone vs. tone freezing: t_(46)_ = 6.014, p < 0.0001; muscimol: pre-tone vs. tone freezing: t_(46)_ = 6.708, p < 0.0001; n_vehicle_ = 13, n_muscimol_ = 13.

GzLMM two-way repeated measures ANOVA of treatment, test stage, and interaction for 21 d context + tone test: χ^2^ treatment _(1)_ = 7.123, p = 0.0076; χ^2^ stage _(1)_ = 41.364, p < 0.0001; χ^2^ interaction _(1)_ = 0.176, p = 0.6751. Sidak’s multiple comparisons test: Pre-tone freezing: vehicle vs. muscimol: t_(26)_ = 2.669, p = 0.0508; Tone freezing: vehicle vs. muscimol: t_(26)_ = 2.683, p = 0.0491; Vehicle: pre-tone vs. tone freezing: t_(26)_ = 7.354, p < 0.0001; muscimol: pre-tone vs. tone freezing: t_(26)_ = 6.431, p < 0.0001; n_vehicle_ = 8, n_muscimol_ = 8.

### Figure 4. Statistics

**E)** GzLMM three-way repeated measures ANOVA of test day, test stage, optogenetic stimulation and interactions for oChIEF stimulated experiment: χ^2^ day _(1)_ = 20.439, p < 0.0001; χ^2^ stage _(1)_ = 100.217, p < 0.0001; χ^2^ stimulation _(1)_ = 5.135, p = 0.0234; χ^2^ interaction(Day×Stage) _(1)_ = 5.805, p = 0.0160; χ^2^ interaction(Day×stimulation) _(1)_ = 2.048, p = 0.15237; χ^2^ interaction(Stage×stimulation) _(1)_ = 2.489, p = 0.1146; χ^2^ interaction(Day×Stage×stimulation) _(1)_ = 4.905, p = 0.0268. Sidak’s multiple comparisons test: Day 7: Pre-tone freezing: Control vs. oChIEF: t_(42)_ = 2.266, p = 0.1601; Day 7: Tone freezing: Control vs. oChIEF: t_(42)_ = 4.454, p = 0.0004; Day 8: Pre-tone freezing: Control vs. oChIEF: t_(42)_ = 3.874, p = 0.0022; Day 8: Tone freezing: Control vs. oChIEF: t_(42)_ = 2.995, p = 0.0272; oChIEF: Tone freezing: Day 7 vs. Day 8: t_(42)_ = 3.810, p = 0.0027; Day 8: oChIEF: Pre-tone vs. Tone freezing: t_(42)_ = 7.283, p < 0.0001; n_Control_ = 7, n_oChIEF_ = 6.

### Figure 5. Statistics

**E)** GzLMM three-way repeated measures ANOVA of test day, test stage, optogenetic inhibition and interactions for CS-US ArchT inhibited experiment: χ^2^ day _(1)_ = 24.559, p < 0.0001; χ^2^ stage _(1)_ = 23.070, p < 0.0001; χ^2^ inhibition _(1)_ = 0.5568, p = 0.4555; χ^2^ interaction(Day×Stage) _(1)_ = 1.659, p = 0.1978; χ^2^ interaction(Day×Inhibition) _(1)_ = 0.294, p = 0.5879; χ^2^ interaction(Stage×Inhibition) _(1)_ = 5.006, p = 0.0253; χ^2^ interaction(Day×Stage×Inhibition) _(1)_ = 1.088, p = 0.2970. Sidak’s multiple comparisons test: Day7: Pre-tone freezing: GFP vs. ArchT: t_(66)_ = 0.746, p = 0.9747; Day 7: Tone freezing: GFP vs. ArchT: t_(66)_ = 3.410, p = 0.0067; Day 8: Pre-tone freezing: GFP vs. ArchT: t_(66)_ = 0.178, p = 1.0000; Day 8: Tone freezing: GFP vs. ArchT: t_(66)_ = 1.145, p = 0.8308; ArchT: Tone freezing: Day 7 vs. Day 8: t_(66)_ = 3.366, p = 0.0076; Day 8: ArchT: Pre-tone vs. Tone freezing: t_(66)_ = 3.475, p = 0.0054; n_GFP_ = 9, n_ArchT_ = 10.

### Supplementary Figure 1. Statistics

GzLMM t test for contextual FC training of 7 d tested group: t_(20)_ = 0.046, p = 0.9635; n_vehicle_ = 11, n_muscimol_ = 12.

### Supplementary Figure 2. Statistics

n_vehicle_ = 6, n_muscimol_ = 6.

**B)** Two-way repeated measures ANOVA of treatment, day, and interaction: F treatment _(1, 10)_ = 0.9320, p = 0.3571; F day _(1, 10)_ = 15.92, p = 0.0026; F interaction _(1, 10)_ = 0.2279, p = 0.6434. Sidak’s multiple comparisons test: Day 1: vehicle vs. muscimol: t = 0.9534, p = 0.5798; Day 2: vehicle vs. muscimol: t = 0.1807, p = 0.9800.

Two-way repeated measures ANOVA of treatment, day, and interaction: F treatment _(1, 10)_ = 1.670, p = 0.2253; F day _(1, 10)_ = 21.48, p = 0.0009; F interaction _(1, 10)_ = 0.8727, p = 0.3722. Sidak’s multiple comparisons test: First day: vehicle vs. muscimol: t = 1.512, p = 0.2708; Second day: vehicle vs. muscimol: t = 0.005307, p > 0.9999.

Two-way repeated measures ANOVA of treatment, day, and interaction: F treatment _(1, 10)_ = 0.01082, p = 0.9192; F day _(1, 10)_ = 0.0003566, p = 0.9853; F interaction _(1, 10)_ = 0.05192, p = 0.8244. Sidak’s multiple comparisons test: Day 1: vehicle vs. muscimol: t = 0.1082, p = 0.9928; Day 2: vehicle vs. muscimol: t = 0.2416, p = 0.9645.

### Supplementary Figure 3. Statistics

GzLMM two-way repeated measures ANOVA of treatment, training stage, and interaction for cued FC training of 7 d tested group: χ^2^ treatment _(1)_ = 0.185, p = 0.6668; χ^2^ stage _(1)_ = 6.308, p = 0.0120; χ^2^ interaction _(1)_ = 0.028, p = 0.8661. Sidak’s multiple comparisons test: Post-shock freezing: vehicle vs. muscimol: t_(28)_ = 0.430, p = 0.8912; Tone freezing: vehicle vs. muscimol: t_(28)_ = 0.297, p = 0.9463; n_vehicle_ = 8, n_muscimol_ = 9.

### Supplementary Figure 4. Statistics

**Aii)** GzLMM t test for contextual FC training of 24 h tested group: t_(16)_ = 1.217, p = 0.2414. n_vehicle_ = 9, n_muscimol_ = 10.

**Aiii)** GzLMM t test for 24 hours context test: t_(16)_ = 3.964, p = 0.0011; n_vehicle_ = 9, n_muscimol_ = 10.

Bii): GzLMM two-way repeated measures ANOVA of treatment, training stage, and interaction for cued FC training of 24 h tested group: χ^2^ treatment _(1)_ = 3.796, p = 0.0514; χ^2^ stage _(1)_ = 0.550, p = 0.4584; χ^2^ interaction _(1)_ = 0.852, p = 0.3560. Sidak’s multiple comparisons test: Post-shock freezing: vehicle vs. muscimol: t_(28)_ = 1.948, p = 0.1192; Tone freezing: vehicle vs. muscimol: t_(28)_ = 1.245, p = 0.3972. n_vehicle_ = 9, n_muscimol_ = 8.

**Biii)** GzLMM Two-way repeated measures ANOVA of treatment, test stage, and interaction for 24 hours cued test: χ^2^ treatment _(1)_ = 0.024, p = 0.8781; χ^2^ stage _(1)_ = 0.108, p = 0.7428; χ^2^ interaction _(1)_ = 12.447, p = 0.0004. Sidak’s multiple comparisons test: Pre-tone freezing: vehicle vs. muscimol: t_(28)_ = 0.153, p = 0.9854; Tone freezing: vehicle vs. muscimol: t_(28)_ = 4.441, p = 0.0003; n_vehicle_ = 9, n_muscimol_ = 8.

### Supplementary Figure 5. Statistics

B, Top (Dorsal) GzLMM Two-way repeated measures ANOVA of treatment, test stage, and interaction for dorsal experiment: χ^2^ treatment _(1)_ = 1.7632, p = 0.1842; χ^2^ stage _(1)_ = 102.565, p < 0.0001; χ^2^ interaction _(1)_ = 0.133, p = 0.7153. Sidak’s multiple comparisons test: Pre-tone freezing: vehicle vs. muscimol: t_(26)_ = 1.328, p = 0.3532; Tone freezing: vehicle vs. muscimol: t_(26)_ = 1.916, p = 0.1283; n_vehicle_ = 8, n_muscimol_ = 8.

B, Middle (Lateral) GzLMM Two-way repeated measures ANOVA of treatment, test stage, and interaction for lateral experiment: χ^2^ treatment _(1)_ = 0.326, p = 0.5680; χ^2^ stage _(1)_ = 27.113, p < 0.0001; χ^2^ interaction _(1)_ < 0.0001, p = 0.9980. Sidak’s multiple comparisons test: Pre-tone freezing: vehicle vs. muscimol: t_(26)_ = 0.571, p = 0.8176; Tone freezing: vehicle vs. muscimol: t_(26)_ = 0.605, p = 0.7978; n_vehicle_ = 8, n_muscimol_ = 8.

B, Bottom (Ventral) GzLMM Two-way repeated measures ANOVA of treatment, test stage, and interaction for ventral experiment: χ^2^ treatment _(1)_ = 0.525, p = 0.4689; χ^2^ stage _(1)_ = 165.9408, p < 0.0001; χ^2^ interaction _(1)_ = 0.259, p = 0.6109. Sidak’s multiple comparisons test: Pre-tone freezing: vehicle vs. muscimol: t_(32)_ = 0.724, p = 0.7235; Tone freezing: vehicle vs. muscimol: t_(32)_ = 0.360, p = 0.9224; n_vehicle_ = 10, n_muscimol_ = 9.

### Supplementary Figure 6. Statistics

B, Left (Cued FC) GzLMM Two-way repeated measures ANOVA of treatment, test stage, and interaction for miss cued test: χ^2^ treatment _(2)_ = 2.5079, p = 0.2854; χ^2^ stage _(1)_ = 85.1604, p < 0.0001; χ^2^ interaction _(2)_ = 33.459, p < 0.0001. Sidak’s multiple comparisons test: Pre-tone freezing: vehicle vs. muscimol: t_(40)_ = 1.457, p = 0.6306; Pre-tone freezing: vehicle vs. miss: t_(40)_ = 1.228, p = 0.7863; Pre-tone freezing: muscimol vs. miss: t_(40)_ = 0.168, p = 1.0000; Tone freezing: vehicle vs. muscimol: t_(40)_ = 5.191, p < 0.0001; Tone freezing: vehicle vs. miss: t_(40)_ = 1.550, p = 0.5636; Tone freezing: muscimol vs. miss: t_(40)_ = 3.559, p = 0.0058; n_vehicle_ = 8, n_muscimol_ = 9, n_miss_ = 7.

### Supplementary Figure 7. Statistics

**B)** GzLMM two-way repeated measures ANOVA of treatment, training stage, and interaction for training of OF-inhibited animals: χ^2^ treatment _(1)_ = 0.685, p = 0.4077; χ^2^ stage _(1)_ = 0.050, p = 0.8231; χ^2^ interaction _(1)_ = 7.700, p = 0.0055. Sidak’s multiple comparisons test: Post-shock freezing: vehicle vs. muscimol: t_(18)_ = 0.828, p = 0.6620; Tone freezing: vehicle vs. muscimol: t_(18)_ = 1.265, p = 0.3945. n_vehicle_ = 6, n_muscimol_ = 6.

GzLMM two-way repeated measures ANOVA of treatment, test stage, and interaction for cued test of OF-inhibited animals: χ^2^ treatment _(1)_ = 0.0697, p = 0.7918; χ^2^ stage _(1)_ = 15.254, p < 0.0001; χ^2^ interaction _(1)_ = 0.103, p = 0.7478. Sidak’s multiple comparisons test: Pre-tone freezing: vehicle vs. muscimol: t_(18)_ = 0.264, p = 0.9579; Tone freezing: vehicle vs. muscimol: t_(18)_ = 0.190, p = 0.9780; n_vehicle_ = 6, n_muscimol_ = 6.

### Supplementary Figure 8. Statistics

**A)** GzLMM two-way repeated measures ANOVA of treatment, training stage, and interaction for cued training of 7 d context + tone tested group: χ^2^ treatment _(1)_ = 0.037, p = 0.8471; χ^2^ stage _(1)_ = 8.741, p = 0.0031; χ^2^ interaction _(1)_ = 8.674, p = 0.0032. Sidak’s multiple comparisons test: Post-shock freezing: vehicle vs. muscimol: t_(46)_ = 0.193, p = 0.9769; Tone freezing: vehicle vs. muscimol: t_(46)_ = 2.216, p = 0.0624. n_vehicle_ = 13, n_muscimol_ = 13.

**B)** GzLMM two-way repeated measures ANOVA of treatment, training stage, and interaction for cued training of 21 d context + tone tested group: χ^2^ treatment _(1)_ = 3.723, p = 0.0537; χ^2^ stage _(1)_ = 0.023, p = 0.8802; χ^2^ interaction _(1)_ = 4.535, p = 0.0332. Sidak’s multiple comparisons test: Post-shock freezing: vehicle vs. muscimol: t_(26)_ = 1.930, p = 0.1251; Tone freezing: vehicle vs. muscimol: t_(26)_ = 0.521, p = 0.8454. n_vehicle_ = 8, n_muscimol_ = 8.

Stage factor refers to freezing between shock and tone, or during tone (post-shock or tone respectively).

### Supplementary Figure 10. Statistics

GzLMM two-way repeated measures ANOVA of optogenetic disruption, training stage, and interaction for oChIEF stimulated group: χ^2^ disruption _(1)_ = 0.007, p = 0.9358; χ^2^ stage _(1)_ = 0.018, p = 0.8946; χ^2^ interaction _(1)_ = 0.0650, p = 0.7988. Sidak’s multiple comparisons test: Post-shock freezing: Control vs. oChIEF: t_(20)_ = 0.081, p = 0.9960; Tone freezing: Control vs. oChIEF: t_(20)_ = 0.113, p = 0.9921. n_Control_ = 7, n_oChIEF_ = 6. Stage factor refers to freezing between shock and tone, or during tone (post-shock or tone respectively).

### Supplementary Figure 12. Statistics

GzLMM two-way repeated measures ANOVA of optogenetic inhibition, training stage, and interaction for cued training of CS-US ArchT inhibited group: χ^2^ inhibition _(1)_ = 0.002, p = 0.9631; χ^2^ stage _(1)_ = 3.173, p = 0.0749; χ^2^ interaction _(1)_ = 5.000, p = 0.0254. Sidak’s multiple comparisons test: Post-shock freezing: GFP vs. ArchT: t_(32)_ = 0.046, p = 0.9987; Tone freezing: GFP vs. ArchT: t_(32)_ = 2.098, p = 0.0859. n_GFP_ = 9, n_ArchT_ = 10. Stage factor refers to freezing between shock and tone, or during tone (post-shock or tone respectively).

### Supplementary Figure 13. Statistics

GzLMM three-way repeated measures ANOVA of test day, test stage, optogenetic inhibition and interactions for inter stimulus ArchT inhibited experiment: χ^2^ day _(1)_ = 20.696, p < 0.0001; χ^2^ stage _(1)_ = 13.802, p = 0.0002; χ^2^ inhibition _(1)_ = 0.424, p = 0.5149; χ^2^ interaction(Day×Stage) _(1)_ = 7.247, p = 0.0071; χ^2^ interaction(Day×Inhibition) _(1)_ = 0.615, p = 0.4330; χ^2^ interaction(Stage×Inhibition) _(1)_ < 0.001, p = 0.9825; χ^2^ interaction(Day×Stage×Inhibition) _(1)_ = 0.294, p = 0.5876. Sidak’s multiple comparisons test: Day7: Pre-tone freezing: GFP vs. ArchT: t_(46)_ = 0.651, p = 0.9875; Day 7: Tone freezing: GFP vs. ArchT: t_(46)_ = 0.704, p = 0.9813; Day 8: Pre-tone freezing: GFP vs. ArchT: t_(46)_ = 0.456, p = 0.9982; Day 8: Tone freezing: GFP vs. ArchT: t_(46)_ = 0.658, p = 0.9868; ArchT: Tone freezing: Day 7 vs. Day 8: t_(46)_ = 0.937, p = 0.9271; Day 8: ArchT: Pre-tone vs. Tone freezing: t_(46)_ = 0.046, p = 1.0000; n_GFP_ = 7, n_ArchT_ = 7.

## Notes

### Competing Interest Statement

The authors have declared no competing interest.

### Summary of Updates

Abstract, introduction results and discussion were modified respect to previous version.

